# High Gamma Cortical Processing of Continuous Speech in Younger and Older Listeners

**DOI:** 10.1101/2019.12.19.883314

**Authors:** Joshua P. Kulasingham, Christian Brodbeck, Alessandro Presacco, Stefanie E. Kuchinsky, Samira Anderson, Jonathan Z. Simon

## Abstract

Neural processing along the ascending auditory pathway is often associated with a progressive reduction in characteristic processing rates. For instance, the well-known frequency-following response (FFR) of the auditory midbrain, as measured with electroencephalography (EEG), is dominated by frequencies from ∼100 Hz to several hundred Hz, phase-locking to the acoustic stimulus at those frequencies. In contrast, cortical responses, whether measured by EEG or magnetoencephalography (MEG), are typically characterized by frequencies of a few Hz to a few tens of Hz, time-locking to acoustic envelope features. In this study we investigated a crossover case, cortically generated responses time-locked to continuous speech features at FFR-like rates. Using MEG, we analyzed responses in the high gamma range of 70–200 Hz to continuous speech using neural source-localized reverse correlation and the corresponding temporal response functions (TRFs). Continuous speech stimuli were presented to 40 subjects (17 younger, 23 older adults) with clinically normal hearing and their MEG responses were analyzed in the 70–200 Hz band. Consistent with the relative insensitivity of MEG to many subcortical structures, the spatiotemporal profile of these response components indicated a cortical origin with ∼40 ms peak latency and a right hemisphere bias. TRF analysis was performed using two separate aspects of the speech stimuli: a) the 70–200 Hz carrier of the speech, and b) the 70–200 Hz temporal modulations in the spectral envelope of the speech stimulus. The response was dominantly driven by the envelope modulation, with a much weaker contribution from the carrier. Age-related differences were also analyzed to investigate a reversal previously seen along the ascending auditory pathway, whereby older listeners show weaker midbrain FFR responses than younger listeners, but, paradoxically, have stronger cortical low frequency responses. In contrast to both these earlier results, this study did not find clear age-related differences in high gamma cortical responses to continuous speech. Cortical responses at FFR-like frequencies shared some properties with midbrain responses at the same frequencies and with cortical responses at much lower frequencies.

**Highlights:**

- Cortical MEG responses time-lock at 80–90 Hz to continuous speech
- Responses primarily driven by high gamma rate fluctuations of the speech envelope
- Response strength and latency are similar for younger and older adults

## 1. Introduction

The human auditory system time-locks to acoustic features of complex sounds, such as speech, as it extracts and encodes relevant information. The characteristic frequency of such time-locked activity is generally thought to decrease along the ascending auditory pathway. For example, subcortical activity at ∼100 Hz and above may directly encode the temporal pitch information of voiced speech (Forte et al., 2017; Krishnan et al., 2004), while cortical activity below ∼10 Hz, which time-locks to the slowly varying envelope of speech, also time-locks to higher level features of language such as phoneme and word boundaries (Brodbeck et al., 2018a). Prior research has also found differences in both subcortical and cortical processing for older and younger listeners (Anderson et al., 2012; Presacco et al., 2016a, 2016b), which suggest age-related auditory temporal processing deficits. These effects have been investigated in human subjects using the complementary non-invasive neural recording techniques of electroencephalography (EEG) and magnetoencephalography (MEG).

The well-known frequency following response (FFR) is one such phase-locked response (Kraus et al., 2017), most commonly measured using EEG, and is believed to originate predominantly from the auditory midbrain (Bidelman, 2015; Smith et al., 1975). The FFR measures the phase-locked response to the fast (∼100 Hz and above), steady state oscillation of a stimulus, such as a repeated speech syllable. The FFR provides insight into the peripheral representation of speech and is a useful tool for investigating temporal processing deficits (Basu et al., 2010; Hornickel et al., 2012; Kraus et al., 2017). In addition, the FFR may be used to investigate the robustness of speech representations in noise or a dual stream paradigm (Yellamsetty and Bidelman, 2019). The FFR is believed to detect the integrated activity of several nonlinear processing stages along the auditory pathway, and hence various nonlinear features of the stimulus can contribute to the FFR (Lerud et al., 2014). Some studies compare and contrast FFRs obtained by averaging or by subtracting responses to stimuli of opposite polarity in order to tease apart these contributions to some extent (Aiken and Picton, 2008; Hornickel et al., 2012).

The neural origins of the FFR have historically been thought to be mainly subcortical areas such as the inferior colliculus (Smith et al., 1975). But recent studies with MEG and EEG have shown that the FFR at ∼100 Hz is not purely generated by subcortical areas, but has contributions from the auditory cortex as well (Bidelman, 2018; Coffey et al., 2017b, 2017a, 2016; Hartmann and Weisz, 2019; Puschmann et al., 2019). Some studies have shown that this cortical contribution is stronger in the right hemisphere (Coffey et al., 2016; Hartmann and Weisz, 2019). The dominantly cortical role in the MEG FFR follows from the reduced sensitivity of gradiometer-based MEG to deep structures such as the auditory midbrain (Baillet, 2017).

However, the repeated speech syllables commonly used to generate the FFR cannot capture the complexities of natural continuous speech. To understand how the brain represents speech in naturalistic environments, cortical low frequency (below ∼10 Hz) responses to continuous speech have been widely studied (Peelle et al., 2013). The MEG and EEG response to continuous speech can be represented using Temporal Response Functions (TRFs) (Ding and Simon, 2012; Lalor et al., 2009) which are linear estimates of time-locked responses to time varying features of the auditory stimulus. The conventional low-frequency TRF time-locks to the slow (below ∼10 Hz) envelope of continuous speech, though the spectrotemporal fine structure of speech can also modulate these cortical low frequency responses (Ding et al., 2014; Ding and Simon, 2012).

Recently, short latency subcortical EEG responses to continuous speech have been found using TRF analysis (Maddox and Lee, 2018), demonstrating that it is possible to detect fast midbrain responses to continuous speech. Early latency responses that phase lock to the fundamental frequency of speech have also been found to be modulated by attention (Forte et al., 2017). One study has also found cortical high gamma MEG responses to speech stimuli, with latencies near 30 ms, that are time-locked to the ∼100 Hz temporal modulation in the envelope of the speech spectrum (up to 2 kHz) (Hertrich et al., 2012). Whether auditory cortex time-locks in the high gamma range to the carrier as well as to the envelope modulation of continuous speech remains unclear.

Further complicating our understanding of the contributions of subcortical and cortical sources to the MEG response is the impact of age-related changes in the auditory pathway (Peelle and Wingfield, 2016). The temporal processing of speech can degrade with age, especially in noisy conditions (Gordon-Salant et al., 2006; He et al., 2008; Hopkins and Moore, 2011). Age-related differences have been found in both the EEG FFR and the MEG low frequency TRF to speech. Older adults have weaker, delayed FFRs with lower phase coherence when compared with younger adults (Anderson et al., 2012; Presacco et al., 2015; Zan et al., 2019). Possible causes include age-related inhibition-excitation imbalance (Caspary et al., 2008) resulting in a loss of temporal precision (Anderson et al., 2012). In a surprising reversal, older adults’ cortex exhibits *exaggerated* low frequency responses (Bidelman et al., 2014; Brodbeck et al., 2018b), even to the point of allowing better stimulus reconstruction via these low frequency cortical responses than in younger adults (Presacco et al., 2016a, 2016b). Several possible explanations, not necessarily exclusive, have been advanced to account for this surprising result, including decrease in inhibition, recruitment of additional brain regions and central compensatory mechanisms (Chambers et al., 2016; Peelle et al., 2010). The fact that fast midbrain responses are reduced with age while slow cortical responses are enhanced might indeed be due to anatomical and physiological differences between midbrain and cortex, but a fair comparison is complicated by the fact that the responses occur at vastly different frequencies. Hence it is entirely unknown whether high gamma cortical responses would show age-related reduction or enhancement.

In this study, we investigated high gamma cortical responses to continuous natural speech using MEG. Unfortunately, MEG responses are known to have relatively poor signal-to-noise ratio (SNR) and decreased power at high gamma frequencies because the cortical sources that dominate MEG responses rarely phase lock in this range at a population level (Lu et al., 2001). In addition, environmental noise and artifacts such as muscular movement can obscure the signal at these higher frequencies (Muthukumaraswamy, 2013). Hence detecting high gamma responses using MEG may require averaging over many trials (as for the FFR), or much longer speech stimuli, to boost SNR. Similarly, detecting subcortical responses to speech may also require high SNR or longer speech stimuli (Maddox and Lee, 2018).

This work investigates whether such high gamma responses can be detected using MEG with a simple experimental paradigm of short duration. MEG recordings of younger and older subjects listening to only six minutes of continuous speech (narration by a male speaker) were investigated using TRF analysis, and such high gamma time-locked responses are indeed found to be present. Just as the low frequency TRF may be compared to a low frequency evoked response, the high gamma TRF may be compared to the FFR in that they both reflect time-locked activity at the stimulus frequency. 70–200 Hz was chosen as the high gamma range because 70 Hz is near the lower end of typical male voice pitch (and well above the 60 Hz of line noise) while 200 Hz is far above most known auditory responses measured by MEG. In addition, source localization was performed to investigate the cortical and subcortical contributions to these high gamma MEG responses. Only six minutes, as opposed to, e.g., 30 minutes, were chosen for the stimulus duration as being typical for an auditory speech experiment that employs multiple stimulus conditions (e.g., several levels of speech in noise). TRF analysis can then be used to investigate time-locked neural processing of a wide variety of stimulus features, from acoustics to semantics (Brodbeck et al., 2018a) simultaneously in the same, short experimental paradigm.

We focused on the following specific research questions. Firstly, are 70–200 Hz MEG responses to continuous speech time-locked to the carrier or to the envelope modulation of the speech spectrum? Unlike FFR analysis, TRF analysis is able to explicitly and simultaneously capture distinct response contributions arising from different stimulus features, in this case from envelope modulation and the carrier, allowing direct comparison of the separate contributions of these features to the response. Secondly, are there any age-related differences in these responses, and if so, do they show age-related decrease, like the EEG FFR, or the opposite, like the cortical low frequency TRFs? Additionally, we investigated if these responses were right lateralized as found in the MEG FFR (Coffey et al., 2016). Such right lateralization would also agree with studies showing right hemispheric dominance for pitch processing in core auditory cortex (Hyde et al., 2008). Finally, we investigated if the responses were influenced by the instantaneous pitch of the speech stimulus.

## 2. Methods

### 2.1 Experiment dataset

The experimental dataset used for this study has been previously described in detail by Presacco et al. (2016a, 2016b), but is here supplemented with eight additional older adults with clinically normal hearing (dataset available online (Kulasingham, 2019a)). The combined dataset consisted of MEG responses recorded from 17 younger adults (age 18–27, mean 22.3, 3 male) and 23 older adults (age 61–78, mean 67.2, 8 male), with clinically normal hearing, while they listened to 60 second portions of an audiobook recording of “The Legend of Sleepy Hollow” by Washington Irving (https://librivox.org/the-legend-of-sleepy-hollow-by-washington-irving). All participants gave informed consent and were paid for their time. Experimental procedures were reviewed and approved by the Institutional Review Board of the University of Maryland. The audio was delivered diotically through 50 Ω sound tubing (E-A-RTONE 3A) attached to E-A-RLINK foam earphones inserted into the ear canal at ∼70 dB sound pressure level via a sound system with flat transfer function from 40 to 3000 Hz. The conditions analyzed in this study consist of two passages of 60 seconds duration presented in quiet (i.e., solo speaker), each of which was repeated three times, for a total of six minutes of MEG data per subject. Subjects were asked beforehand to silently count the number of occurrences of a particular word and report it to the experimenter at the conclusion of each trial, in order to encourage attention to the auditory stimuli. Handedness of the participants was assessed with the Edinburgh handedness scale (Oldfield, 1971), which can range from –1 (complete left-dominance) to 1 (complete right-dominance). To exclude lateralization bias due to handedness, all analyses were performed again excluding the 9 subjects scoring below 0.5. The only qualitative change in the results was a loss of right hemispheric dominance in younger subjects (discussed below).

### 2.2 MEG data collection and preprocessing

MEG data was recorded from a 157 axial gradiometer whole head KIT MEG system while subjects were resting in the supine position in a magnetically shielded room. The data was recorded at a sampling rate of 1 kHz with an online 200 Hz low pass filter with a wide transition band above 200 Hz, and a 60 Hz notch filter. Data was preprocessed in MATLAB by first automatically excluding saturating channels and then applying time-shift principal component analysis (de Cheveigné and Simon, 2007) to remove external noise, and sensor noise suppression (de Cheveigné and Simon, 2008) to suppress channel artifacts. On average, two MEG channels were excluded during these stages. All subsequent analyses were performed in mne-python 0.17.0 (Gramfort, 2013; Gramfort et al., 2014) and eelbrain 0.30 (Brodbeck et al., 2019); code available online (Kulasingham, 2019b). The MEG data was filtered in the band 70–200 Hz (high gamma band) using a FIR filter described below, and six 60 second epochs during which the stimulus was presented were extracted for analysis. The band 70–200 Hz was chosen since the pitch of the male speaker typically falls in this range, and 200 Hz is above most known auditory cortical responses. The data was resampled to 500 Hz for all further analysis.

### 2.3 Stimulus representation

As discussed above, prior work on the FFR has shown that time-locked neural responses are sensitive to both the carrier and the envelope of an auditory stimulus. Similarly, time-locked responses to speech in the high gamma range may be driven either by the high gamma carrier, or by high gamma modulation in the envelope of even higher frequencies. Accordingly, two distinct representations of the speech stimulus were used as predictors for the TRF model (see Fig. 1). For the former case, the carrier predictor was constructed by resampling the speech waveform to 1 kHz (using the mne-python function ‘resample’) and bandpass filtering from 70–200 Hz using the same filter as above. This carrier predictor captures the high gamma rate modulation in the speech waveform itself. For the latter case, the envelope modulation predictor was constructed from the high gamma modulation in the envelope of the highpassed stimulus waveform (envelopes are only well-defined when they modulate carriers of higher frequencies than those of the modulations themselves; Rosen, 1992). Specifically, first the speech was transformed into an auditory spectrogram representation by computing the acoustic energy in the speech waveform for each frequency bin in the range 300–4000 Hz at millisecond resolution using a model of the auditory periphery (Yang et al., 1992). The range 300–4000 Hz was chosen in order to have a clear separation between the upper end of the high gamma range (200 Hz) and because the auditory stimulus was presented through air tubes which attenuate frequencies above 4000 Hz. This auditory spectrogram is a 2-dimensional matrix representation of the acoustic envelope over time for different frequency bins. Each frequency bin component of this spectrogram was then filtered using the same 70–200 Hz bandpass filter as above, producing a 70–200 Hz band limited envelope for each bin. Finally, the resulting 2-dimensional matrix was averaged across frequency bins to provide a single signal, resulting in the envelope modulation predictor. Thus, this predictor captures the 70–200 Hz temporal modulation in the 300–4000 Hz envelope of the speech waveform. These two predictors were resampled to 500 Hz and used for all further TRF analysis. Even though the two predictors are correlated at *r* = −0.42, the TRF analysis is able to separate the neural response to each of them (negative correlations are common for a carrier and the corresponding non-linearly related envelope of another frequency band with different cochlear delays).

**Fig. 1.**
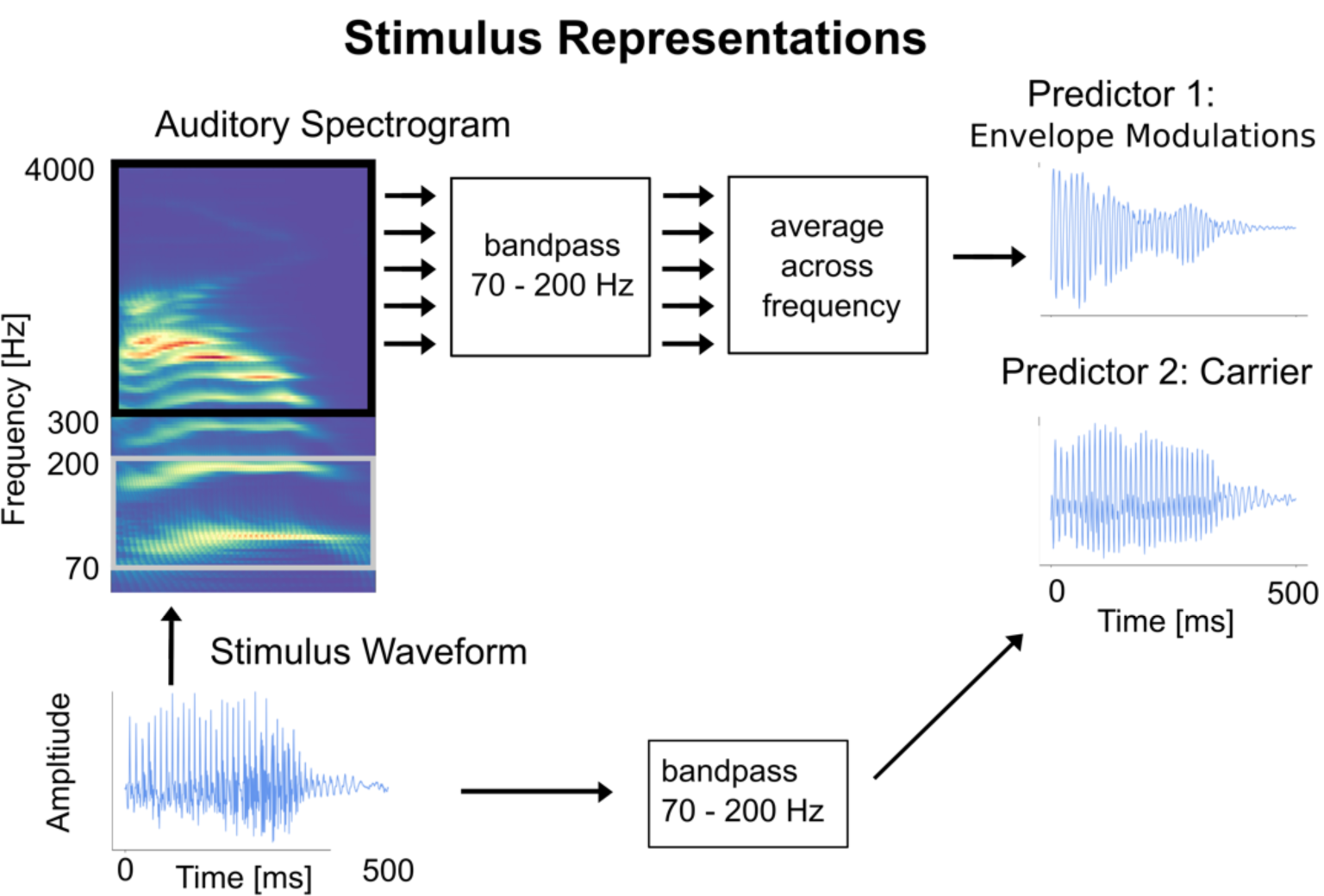
Stimulus Representations. The stimulus waveform for a representative 500 ms speech segment is shown along with its auditory spectrogram and the two predictors: carrier and envelope modulation. The predictors are correlated (Pearson’s *r* = –0.42) but have noticeably distinct waveforms.

The 70–200 Hz bandpass filter was formed using the default FIR filter in mne-python with an upper and lower transition bandwidth of 5 Hz, at 1 kHz sampling frequency, but applied twice in a forward fashion to the data. This resulted in a combined filter of length 1322 with a phase delay of 660 ms. Other bandpass filters were also employed as alternatives, including IIR minimum-phase-delay Bessel filters (results not shown); no results depended critically on the filters used.

### 2.4 Neural source localization

Before each MEG recording, the head shape of each subject was digitized using a Polhemus 3SPACE FASTRAK system, after which five marker coils were attached. The marker coil locations were measured while the subject’s head was positioned in the MEG scanner before and after the experiment, in order to determine the position of the head with respect to the MEG sensors. Source localization was performed using the mne-python software package. The marker coil locations and the digitized head shape were used to coregister the template Freesurfer ‘fsaverage’ brain (Fischl, 2012) using rotation, translation and uniform scaling. A volume source space was formed by dividing the brain volume into a grid of 7 mm sized voxels. This source space was used to compute an inverse operator using minimum norm estimation (MNE) (Gramfort et al., 2014) and dynamical statistical parametric mapping (dSPM) (Dale et al., 2000) with a depth weighting parameter of 0.8, and a noise covariance matrix estimated from empty room data. This method results in a 3-dimensional current dipole vector with magnitude and direction at each voxel. The Freesurfer ‘aparc+aseg’ parcellation was used to define cortical and subcortical regions of interest (ROIs). The cortical ROI consisted of voxels in the gray and white matter of the brain that were closest to the temporal lobe Freesurfer ‘aparc’ parcellations (‘aparc’ labels: ‘transversetemporal’, ‘superiortemporal’, ‘inferiortemporal’, ‘bankssts’). A few additional voxels surrounding auditory cortex (within 20 mm) were included in the ROI solely to ensure that the source localized responses not be misleadingly focal (distributed source localization with MNE has a large spatial spread). The subcortical ROI was selected to consist of voxels that were in the Freesurfer ‘aseg’ ‘Brain-Stem’ segmentation. All brain plots show the maximum intensity projection of the voxels onto a 2-dimensional plane, with an overlaid ‘fsaverage’ brain schematic (implemented in eelbrain). Minimum norm estimation in volume source space may lead to spatial leakage from the true neural source to neighboring voxels. In order to characterize this artifactual spatial leakage, a single current dipole in Heschl’s gyrus was simulated, projected into sensor space, and then projected into volume source space (see Appendix). Additionally, a separate cortical surface source space model was also used; results obtained using this method were not qualitatively different than those of the volume space model (see Appendix).

### 2.5 Temporal response functions

The simplest linear model used to estimate the TRF is given by

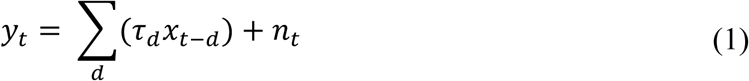

where *y*_*t*_ is the response at a neural source for time *t, x*_*t-d*_ is the time shifted predictor with a time lag of *d, τ*_*d*_ is the TRF value at lag *d* and *n*_*t*_ is the residual noise. The TRF is the set of time-dependent weights, of a linear combination of current and past samples of the predictor, that best predicts the current neural response at that neural source (Lalor et al., 2009). Hence the TRF can also be interpreted as the average time-locked response to a predictor impulse. In this investigation, a TRF model with two predictors, envelope modulation and carrier, was used.

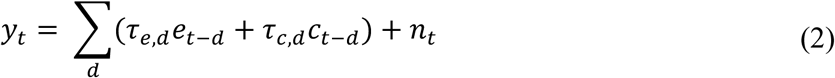

Where *e*_*t−d*_ is the delayed <u>e</u>nvelope modulation predictor and *τ*_*e,d*_ the corresponding envelope modulation TRF, *c*_*t−d*_ is the delayed <u>c</u>arrier predictor and *τ*_*c,d*_ the corresponding carrier TRF. In this model, the two predictors compete against each other to explain response variance, which results in larger TRFs for the predictor that contributes more to the neural response. The model parameters were estimated jointly, such that the model is not affected by the ordering of the predictors. TRF estimation, for lags from –40 to 200 ms, was performed with the boosting algorithm and early stopping based on cross validation (David et al., 2007) as implemented in eelbrain. The boosting algorithm may result in overly sparse TRFs, and hence an overlapping basis of 4 ms Hamming windows (with 1 ms spacing) was used in order to allow smoothly varying responses; altering the Hamming window duration did not substantively affect the results. For the volume source space, the neural response at each voxel is a 3-dimensional current vector. Accordingly, for each voxel, a TRF vector was computed using the boosting algorithm and was used to predict the neural response vector. For each voxel, the prediction accuracy was assessed through the average dot product between the normalized predicted and true response, which varies between −1 and 1 in analogy to the Pearson correlation coefficient.

### 2.6 Pitch analysis

Prior studies have suggested that neural time-locking at high gamma rates may reflect processing of pitch related features of the speech (Smith et al., 1978). In order to investigate the extent to which the response oscillations were influenced by the pitch frequency of the speech stimulus, a simple pitch analysis was performed as follows. The pitch of the speech signal was extracted using Praat (Boersma, 1993; Boersma and Weenick, 2018) in sliding 40 ms windows and used to mark times when the pitch was above or below the median pitch value (98.11 Hz). This algorithm is a better approximation of the percept of pitch than simply dividing the stimulus based on its frequency content, thus allowing subsequent analysis to be done on a neurally relevant feature of the stimulus. Two new ‘high pitch’ predictors were formed based on the previous two predictors (envelope modulation and carrier) by zeroing out the times when the pitch was below the median. Similarly, ‘low pitch’ predictors were formed by zeroing out times when the pitch was above the median. Time windows without a stable pitch estimate were set to zero in all predictors. Hence four new predictors were created: high pitch envelope modulation, low pitch envelope modulation, high pitch carrier and low pitch carrier. All 4 predictors were used simultaneously in a competing TRF model analogous to that in Eq. (2).

### 2.7 Statistical tests

Statistical tests were performed across subjects by comparing the TRF model to a noise model. The predictor was circularly shifted in time and TRFs were estimated using this time-shifted predictors as noise models (Brodbeck et al., 2018a, 2018b). This preserves the local temporal structure of the predictor while removing the temporal relationship between the predictor and the response. Circular shifts of duration 15, 30 and 45 seconds were used to form three noise models. For each voxel, the prediction accuracies of the true model were compared to the average prediction accuracies of the three noise models as a measure of model fit. Since all the predictors in the model are fit jointly, this results in one joint prediction accuracy for all the predictors for each voxel.

To account for variability in neural source locations due to mapping the responses of individual subjects onto the ‘fsaverage’ brain, these coefficients were spatially smoothed using a Gaussian window with 5 mm standard deviation. Nonparametric permutation tests (Nichols and Holmes, 2002) and Threshold Free Cluster Enhancement (TFCE) (Smith and Nichols, 2009) were used to control for multiple comparisons. This method, as outlined in full in Brodbeck et al., 2018c, 2018a, is implemented in eelbrain, and is briefly recounted here. Firstly, a paired sample *t*-value was evaluated for each neural source, across subjects, from the difference of the prediction accuracies of the true model and the average of the three noise models after rescaling using Fisher’s *z*-transform. Then the TFCE algorithm was applied to those *t*-values, which enhanced continuous clusters of large values, based on the assumption that significant neural activity would have a larger spatial spread than spurious noise peaks. This procedure was repeated 10,000 times with random permutations of the data where the labels of the condition were flipped on a randomly selected subset of the subjects. A distribution of TFCE values was formed using the maximum TFCE value of each permutation to correct for multiple comparisons across the brain volume. Any value of the original TFCE map that exceeded the 95th percentile of the distribution was considered as significant at the 5% significance level. This corresponds to a one-tailed test of whether the true model increases the prediction accuracy over the noise model. In cases where both sides of the comparison are important, corresponding two-tailed tests were used (as explained below, for e.g., left vs. right, younger vs. older, envelope vs. carrier). In all subsequent results, the maximum or minimum t-value across voxels is reported as *t*_*max*_ or *t*_*min*_ respectively.

The TRF itself was also tested for significance against the noise model in a similar manner. In the volume source space, a TRF that consists of a 3-dimensional vector which varies with time was estimated for each voxel, representing the estimated current dipole amplitude and direction at that voxel. The amplitudes of these TRF vectors for the true model and the average noise model were used for significance testing. The TRF amplitudes were spatially smoothed using the same Gaussian window before performing the tests. A one-tailed test was done with paired sample *t*-values and TFCE, and the procedure is identical to that outlined previously, with the added dimension of time (Brodbeck et al., 2018a).

Lateralization tests were performed to check for hemispheric asymmetry. The volume source space estimates in the cortical ROI were separated into left and right hemispheres and, as above, the prediction accuracies were spatially smoothed with the same Gaussian window. The prediction accuracies of the average noise model were subtracted from that of the true model and paired sample *t*-values with TFCE in a two-tailed test were used to test for significant differences between each of the corresponding left and right voxels.

Age-related differences were assessed between the younger and older groups. The difference of prediction accuracies between the true TRF model and the average of the noise TRF models were used to form independent sample *t*-values for each source across age groups after which a two-tailed test was performed with TFCE. Significant differences in lateralization across age groups were assessed by subtracting the prediction accuracies of the left hemisphere from the right hemisphere and then conducting independent samples tests across age groups as described above. The peak latency of the TRFs was also tested for significant differences across age groups. The latency of the maximum value of the norm of the TRF vectors in the time range of significant responses (20–70 ms) was used to test for peak latency differences across age groups using a two-tailed test with independent sample *t*-values and TFCE.

To further investigate differences by age across both low frequency and high frequency (i.e., high gamma) responses, two additional models were analyzed; a low frequency (1–10 Hz) TRF and a high frequency TRF with the same parameters as the above models, but using cortical surface source space. An ANOVA was performed on the prediction accuracies of these two models with factors TRF frequency (high or low) and age (young or old) (detailed methods and results in Appendix).

## 3. Results

### 3.1 Cortical origins of high gamma responses to continuous speech

Per-voxel TRFs in volume source space were estimated in the high gamma range for the two ROIs: the temporal lobes, and the brainstem (plus its surrounding volume). The prediction accuracies of the competing stimulus model described above for high-frequency responses (mean = 0.021, std = 0.003) were much smaller (factor of 3) than those resulting from low frequency cortical TRFs (Brodbeck et al., 2018a), indicating that these responses are weaker than slow cortical responses. This is not surprising, as the spectral power of the MEG response decays with frequency. Noise floor models, used to test for significant responses, generated corresponding noise model prediction accuracies (mean = 0.018, std = 0.001). For each voxel, a one-tailed test with paired sample *t*-values and TFCE (to account for multiple comparisons) was used to test for significant increases in the prediction accuracies of the true model against the noise model across subjects. A large portion of the voxels showed a significant increase in prediction accuracy (younger subjects *t*_*max*_ = 6.19, *p* < 0.001; older subjects *t*_*max*_ = 5.66, *p* < 0.001; see Fig. 2A). The disproportionate extent of this result is not unexpected, however, due to the large spatial spread of MNE volume source space estimates. The prediction accuracy over the noise model for voxels in Heschl’s gyrus was significantly larger than that within the subcortical ROI for both age groups (two-tailed paired sample *t*-test; younger subjects *t* = 3.67, *p* = 0.002; older subjects *t* = 2.65, *p* = 0.015; difference across age not significant). Although some voxels in the subcortical ROI are significant, this can be ascribed to artifactual leakage arising from the source localization algorithm (see simulation in Appendix).

**Fig. 2.**
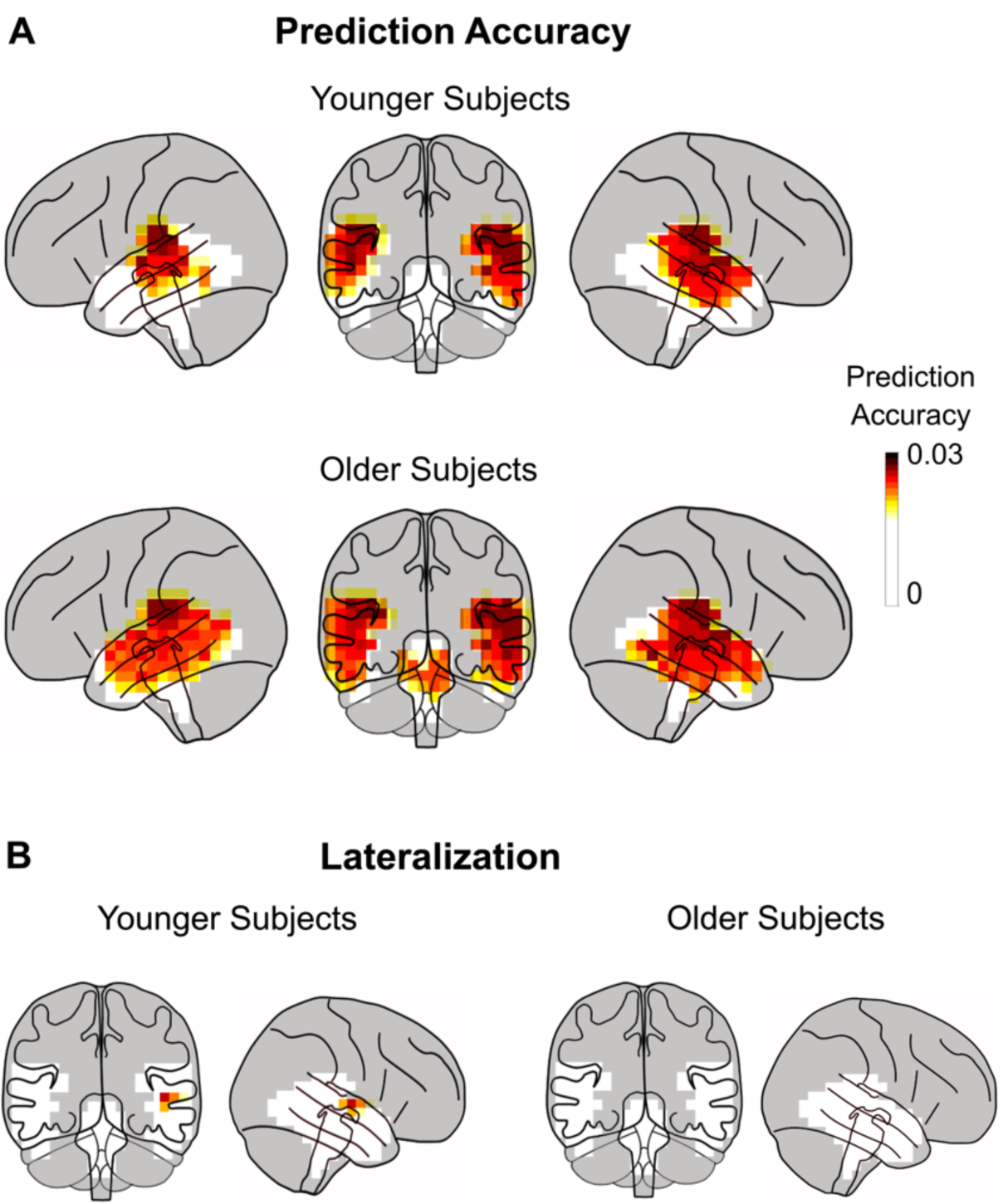
Prediction Accuracy of Volume Source Localized TRFs. **A**. Prediction accuracy using the TRF model for each voxel in the volume source space ROIs (non-gray regions) averaged across subjects. Only ROI voxels for which model prediction accuracy significantly increased over the noise model are plotted (*p* < 0.05, corrected). The prediction accuracy is larger in cortical areas than in subcortical areas. Plots are of the maximum intensity projection, with an overlay of the brain. When taking into account expected MEG volume source localization leakage, these results are consistent with the response originating solely from cortical areas and with a right hemispheric bias. **B**. An area in the right hemisphere near the auditory cortex is significantly more predictive than the left hemisphere, but only in the younger subjects.

Lateralization differences were tested using the prediction accuracy at each voxel. The prediction accuracy of the average noise model was subtracted from that of the true model and a two-tailed test with paired sample *t*-values and TFCE was performed for significant differences in the left and right hemispheres. The tests revealed significantly higher prediction accuracies for younger subjects in the right hemisphere than in the left (*t*_*max*_ = 3.81, *p* = 0.035), but only for a few voxels (1.6%) in the temporal lobe close to auditory areas (see Fig. 2B). No significant differences in lateralization were seen for older subjects (*t*_*max*_ = 3.41, *t*_*min*_ = –1.52, *p* > 0.09), nor was lateralization significantly different across age groups (independent samples test; *t*_*max*_ = 1.93, *t*_*min*_ = –2.28, *p* > 0.88). When the analysis was constrained to only right-handed subjects (13 younger, 18 older; see Methods for details), the only resulting change was that no voxels were significantly right lateralized in either age group.

The TRFs at each source voxel are represented by a 3-dimensional current vector that varies over the time lags. Hence for each voxel and time lag, the amplitude of the TRF vector for the true model was tested for significance against the average of the noise models across subjects using a one-tailed test with paired sample *t*-values and TFCE. The TRFs for the envelope modulation predictor in the cortical ROI were significant (younger *t*_*max*_ = 5.38, *p* < 0.001; older *t*_*max*_ = 4.69, *p* < 0.001) starting at a time lag of 23 ms, and ending at 63 ms, with an average peak latency of 40 ms (see Fig. 3A). The TRF current dipoles oscillate with alternating direction between successive amplitude peaks. However, in all subsequent TRF plots, the TRF amplitude is shown, and not signed current values, and hence signal troughs and peaks both appear as peaks. The subcortical ROI was also analyzed in a similar manner and the TRF showed significance in a much smaller time range of 31–35 ms only for older subjects (younger *t*_*max*_ = 2.96, *p* > 0.13; older *t*_*max*_ = 3.69, *p* < 0.01) (see Fig 3B). There was no significant difference in amplitudes between younger and older subjects (cortical ROI *t*_*max*_ = 3.7, *t*_*min*_ = –3.38, *p* > 0.18; subcortical ROI *t*_*max*_ = 3.05, *t*_*min*_ = –3.39, *p* > 0.45). The TRF responses oscillate at a frequency of ∼80 Hz (see below for a more detailed spectral analysis). The amplitude of these TRFs was significantly larger in voxels in Heschl’s gyrus than in the subcortical ROI (two-tailed test with paired sample t-values on the *l*2 norm of the TRFs across subjects: younger *t* = 3.51, *p* = 0.003; older *t* = 4.52, *p* < 0.001). Since the subcortical TRFs also have a similar latency and shape to the cortical TRFs, and because a latency of 23 to 63 ms is late for a subcortical response, these subcortical TRFs are consistent with artifactual leakage from the cortical TRFs due to the spatial spread of MNE source localization. Simulated volume source estimates for current dipoles originating only in Heschl’s gyrus generated a spatial distribution of TRF directions consistent with the experimental data (see Appendix), i.e. the spatial spread of MNE localized cortical responses resulted in apparent TRF vectors even in the subcortical ROI. These results indicate that the response originates predominantly from cortical regions.

**Fig. 3.**
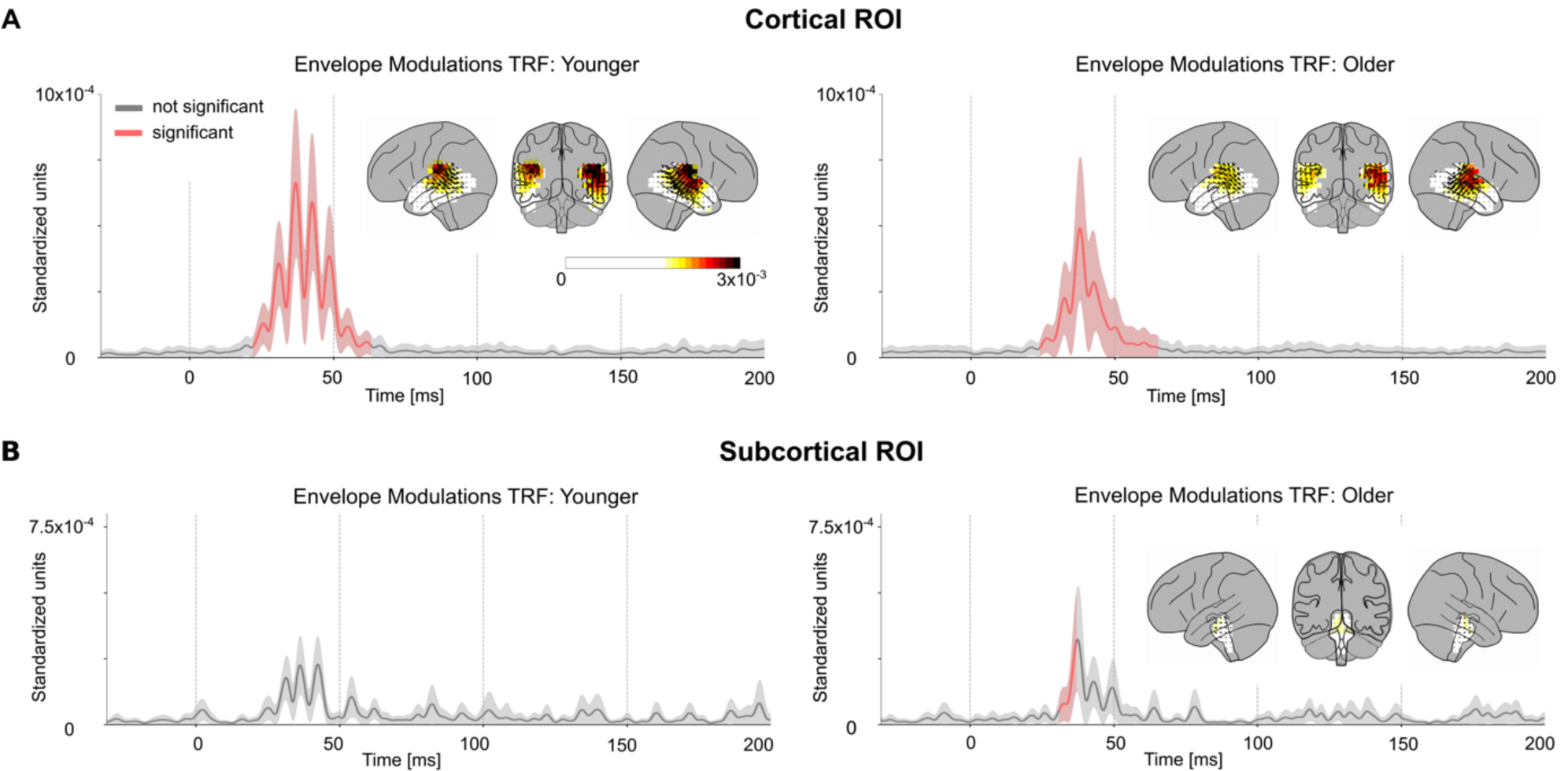
Volume Source Localized Envelope Modulation TRFs. The amplitude of the TRF vectors for the envelope modulation predictor averaged across voxels in the ROI, and, the mean ± (standard error) across subjects is plotted in the cortical (A) and subcortical (B) ROIs. Red curves are time points when the TRF showed a significant increase in amplitude over noise. The TRF was resampled to 1000 Hz for visualization purposes. The TRF shows a clear response with a peak latency of ∼40 ms. The distribution of TRF vectors in the brain at each voxel at the time with the maximum response are plotted as an inset for each TRF, with color representing response strength and the arrows representing the TRF directions. The color bar represents the response strength for all 4 brain insets. The response oscillates around a frequency of ∼80 Hz and is much stronger in the cortical ROI compared to the subcortical ROI. Note that since only the TRF amplitude is shown, and not signed current values, signal troughs and peaks both appear as peaks. In the original, signed TRFs, the current direction alternates between successive amplitude peaks. The latency and amplitude of the response suggests a predominantly cortical origin.

### 3.2 Responses to the envelope modulation and the carrier

Next, the neural response to the carrier was compared with that to the envelope modulation. The carrier TRF was also tested for significance using a corresponding noise model (as employed above). The carrier TRF showed weak responses that were only significant in the cortical ROI between 33–51 ms (younger *t*_*max*_ = 3.70, *p* = 0.042; older *t*_*max*_ = 4.7, *p* < 0.001) (see Fig. 4A, B). Although the carrier and envelope modulation predictors are correlated (*r* = –0.42), the TRF analysis is able to separate the contributions of these two predictors remarkably well. Two-tailed paired sample *t*-values and TFCE were used to test for a significant increase of the *l*2 norm of the envelope modulation TRF when compared to the carrier TRF in a time window of 20–70 ms in the cortical ROI (see Fig. 5A). This test was significant for both younger (*t*_*max*_ = 4.38, *p* = 0.002) and older (*t*_*max*_ = 3.63, *p* = 0.017) subjects. However, this test did not find a significant increase in the envelope modulation TRF over the carrier TRF in the subcortical ROI for either younger (*t*_*max*_ =0.045, *p* > 0.32) or older subjects (*t*_*max*_ = 0.89, *p* > 0.36). Since the TRF analysis allows both stimulus predictors to directly compete for explaining response variance, the results strongly indicate that the response is primarily due to the envelope modulation over the carrier.

**Fig. 4.**
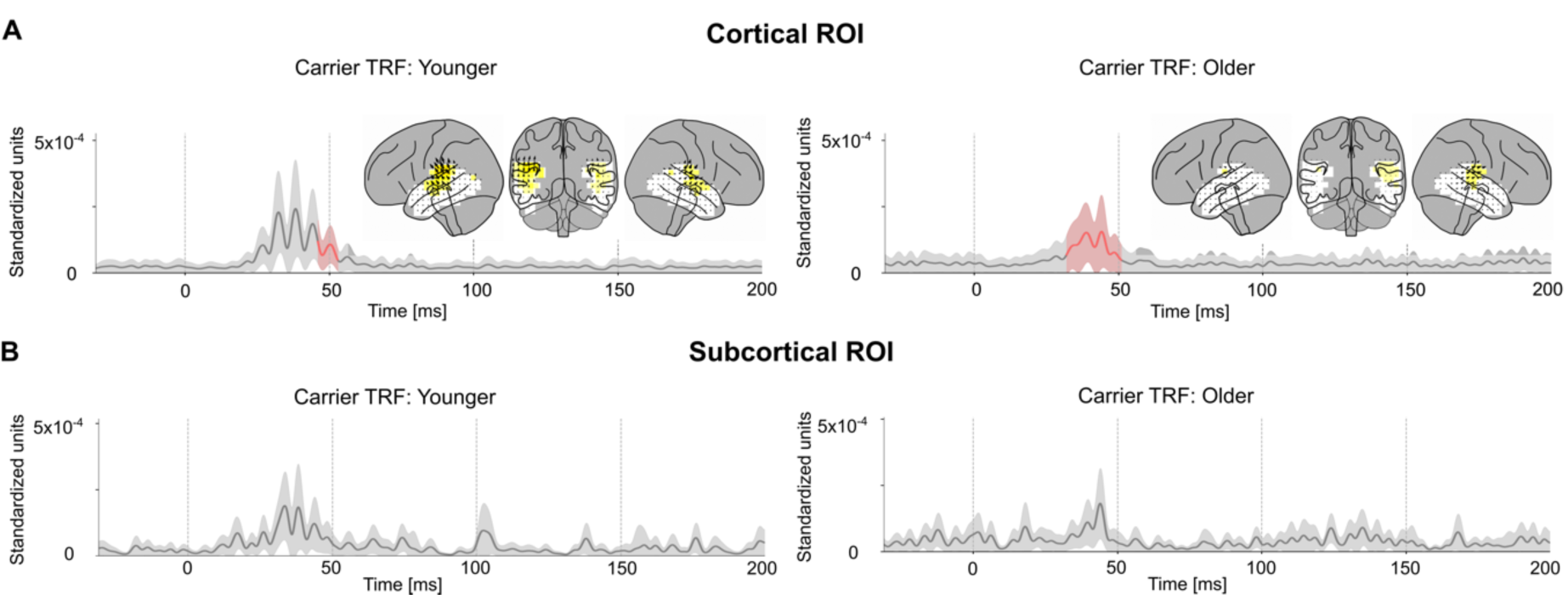
Volume Source Localized Carrier TRFs. The amplitude of the TRF vectors for the carrier predictor averaged across sources. Mean ± (standard error) across subjects is shown, analogous to Fig. 3. For comparison, the axis and color scale are identical to that in Fig. 3. The TRF shows a weaker response compared to the case of envelope modulation, with a peak latency of ∼40 ms, that is significant in the cortical ROI for both groups, and over a longer time interval for older subjects. Comparison with Fig. 3 suggests that the high gamma response is dominated by the envelope modulation over the carrier.

**Fig. 5.**
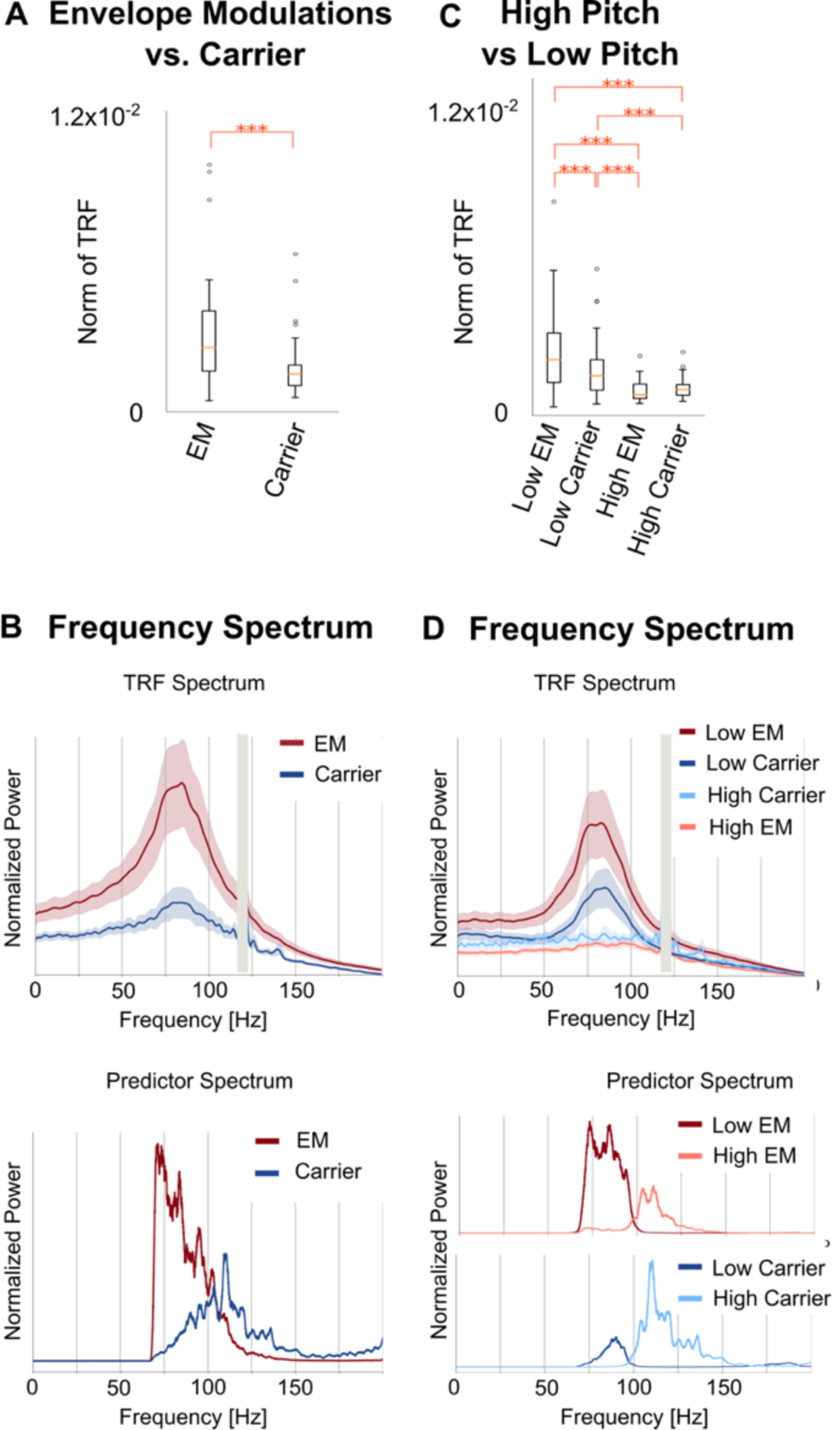
Comparison of Responses to the Envelope Modulation and to the Carrier. **A**. The norm of the TRF between 20 ms and 70 ms was larger in the envelope modulation TRF than the carrier TRF (*** *p* < 0.001). Boxplots after combining both age groups are shown. **B**. The frequency spectrum of the TRF reveals that the oscillation has a broad peak around 80 Hz (vertical gray bars denote a narrow frequency band excluded from analysis because of 120 Hz line noise). In contrast, the predictors’ peaks are displaced in frequency from the TRF peak, either well below (for the envelope modulation) or well above (for the carrier). Note that the sharp cutoff in the envelope modulation spectrum at 70 Hz arises from the bandpass filter used in analysis: without the bandpass filter the spectrum would continue rising toward lower frequencies. **C**. The norm of the TRF for the pitch-separated model between 20 ms and 70 ms was larger in the low pitch TRFs than the high pitch TRFs for both envelope modulation and carrier (*** p < 0.001). **D**. The frequency spectrum of the low-pitch TRF has a peak around 80 Hz, while the high pitch TRF does not show any peaks. This suggests that the TRF is dominantly driven by the low-pitch segments of the speech waveform. The spectra of the corresponding high and low pitch predictors are also shown, highlighting the clear separation of the spectra at the median pitch frequency of 98 Hz.

### 3.3 Age-related differences

Statistical tests were performed for age-related differences between older and younger subjects on both the prediction accuracy and the TRFs. Two-tailed tests of prediction accuracy with independent sample *t*-values and TFCE indicated no significant difference (cortical ROI *t*_*max*_ = 1.17, *t*_*min*_ = –2.72, *p* > 0.44; subcortical ROI *t*_*max*_ = –0.78, *t*_*min*_ = –1.37, *p* > 0.38). Similarly, no voxels or time points were significantly different in either the envelope modulation TRF (cortical ROI *t*_*max*_ = 3.7, *t*_*min*_ = –3.38, *p* > 0.18; subcortical ROI *t*_*max*_ = 3.05, *t*_*min*_ = –3.39, *p* > 0.45) or the carrier TRF (cortical ROI *t*_*max*_ = 3.34, *t*_*min*_ = –3.89, *p* > 0.25; subcortical ROI *t*_*max*_ = 2.69, *t*_*min*_ = – 3.10, *p* > 0.18). In addition, the cortical ROI TRFs showed no significant differences across age groups in peak latency (envelope modulation TRF *t*_*max*_ = 1.82, *t*_*min*_ = –2.62, *p* > 0.5; carrier TRF *t*_*max*_ = 2.79, *t*_*min*_ = –2.32, *p* > 0.53). An additional analysis was performed using surface source space TRFs as described in detail in the Appendix. Both high (70–200 Hz) and low (1–10 Hz) frequency TRFs were computed in surface source space, and model prediction accuracy was assessed with an ANOVA with factors TRF frequency and age. The ANOVA showed a significant frequency × age interaction (*F*_*1,38*_ = 6.46, *p* = 0.015), suggesting that age related differences are indeed not consistent across high and low frequency responses (detailed results in Appendix), i.e. present at low but not at high frequencies.

### 3.4 Pitch analysis

To further understand the contributions of these predictors to the TRF oscillations, the frequency spectrum of the TRFs and the predictors were compared (see Fig. 5B). The frequency spectrum of the average TRFs showed a broad peak centered near 80 Hz for both predictors and both age groups (envelope TRF spectral peak mean = 81 Hz, std = 5 Hz; carrier TRF spectral peak mean = 82 Hz, std = 8 Hz). In contrast, the spectral peak of the predictor variables was near 110–120 Hz for the carrier, and near 70–75 Hz for the envelope modulation. Since the TRF peak frequency did not match the peak power in either of the predictors, a further analysis was performed after separating the stimulus into high- and low-pitch time segments (see Methods). This resulted in a model with 4 predictors and their corresponding TRFs: high/low-pitch envelope modulation and high/low-pitch carrier. The low-pitch envelope modulation TRFs and low-pitch carrier TRFs are broadly similar to those of the earlier analysis (see Fig. 6). These TRFs show more significant regions than the previous analysis, although the two models (one with 2 predictors, the other with 4 predictors) cannot be directly compared since an increased number of predictors has more degrees of freedom and allows for the model to predict more of the signal. The TRF amplitudes were significantly larger in the low pitch TRFs when compared to the high pitch TRFs (see Fig. 5C; envelope modulation *t*_*max*_ = 7.6, *p* < 0.001; carrier *t*_*max*_ = 3.78, *p* = 0.013). In addition, the spectra of the low pitch TRFs peak near 80 Hz similar to the low pitch predictors (envelope TRF spectral peak mean = 81 Hz, std = 6 Hz; carrier TRF spectral peak mean = 82 Hz, std = 4 Hz), while the high pitch TRFs do not have a clear peak (see Fig. 5D). This suggests that the TRF oscillation is driven mainly by the segments of the stimulus with pitch below 100 Hz, and that responses to stimulus pitches above 100 Hz are not easily detected by this analysis.

**Fig. 6.**
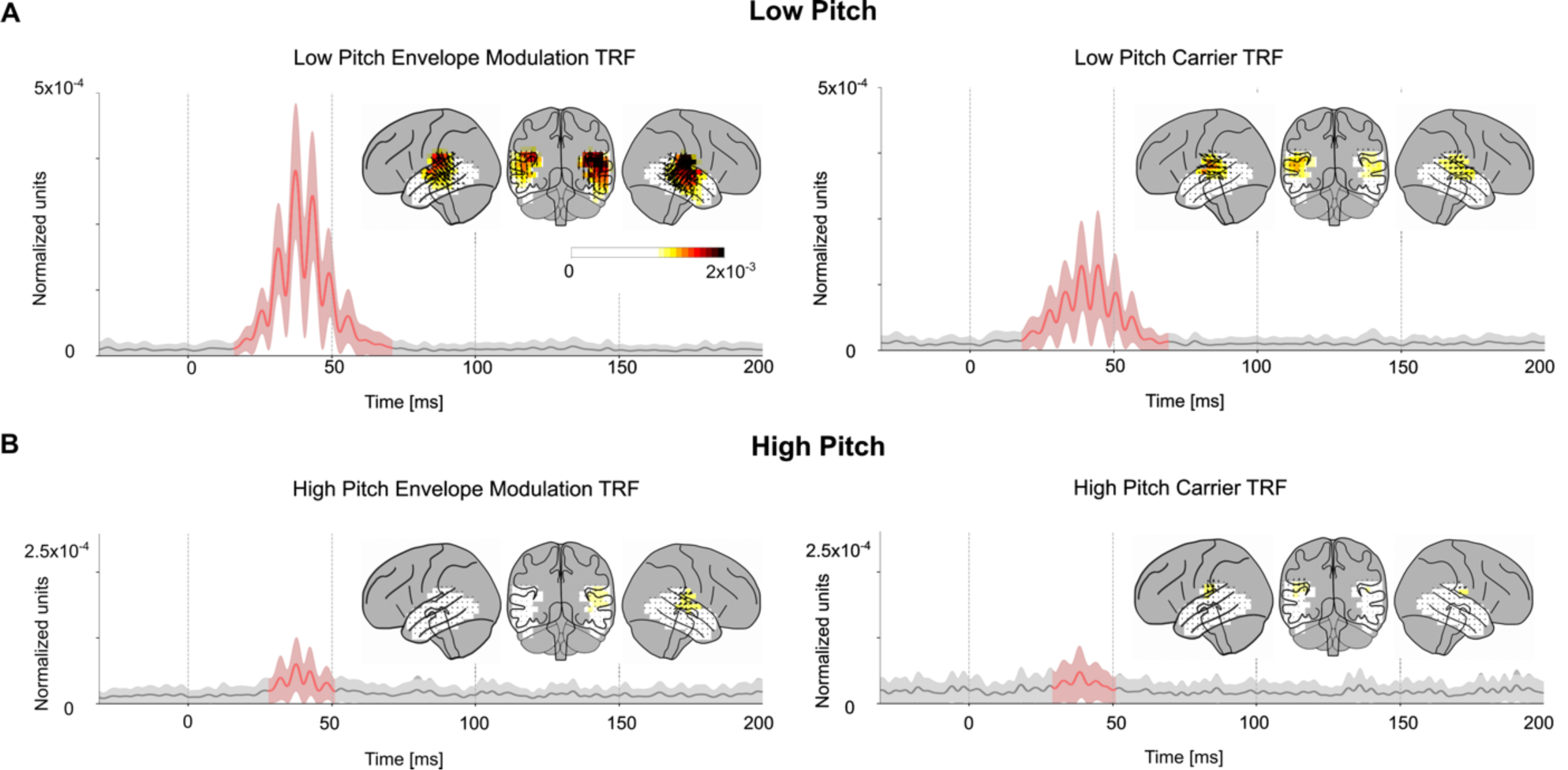
Pitch-separated TRFs. The amplitude of the low pitch TRF (A) and high pitch TRF (B) vectors for both the envelope modulation and carrier predictors averaged across sources. Mean ± (standard error) across subjects is shown, analogous to Figs. 3, 4. The axis and color scale are smaller than those in Fig. 3, 4, since the pitch separated TRFs are each based on a subset of the stimulus predictors, and hence have weaker amplitudes. All four TRFs show significant regions around 40 ms, but the low pitch envelope modulation TRF is the strongest, followed by the low pitch carrier TRF. This indicates that the high gamma response is time-locked to the low pitch segments of the speech stimulus.

## 4. Discussion

In this study, we investigated high gamma time-locked responses to continuous speech measured using MEG. Such responses were found, and their volume source localized TRFs provided evidence that these responses originated from cortical areas with a peak response latency of approximately 40 ms. The responses showed a significant right hemispheric asymmetry. These responses oscillate with a frequency of approximately 80 Hz and track the low pitch segments of the speech stimulus. We also showed that the response is significantly stronger to the envelope modulation than the carrier. Surprisingly, there were no significant age-related differences in response amplitude, latency, localization or predictive power. This is in contrast to age-related differences seen in both the subcortical EEG FFR (younger > older) and the cortical low frequency TRF (older > younger).

### 4.1 MEG sensitivity to high gamma responses

MEG signals are known to have poor SNR at high frequencies (≳100 Hz) (Hansen et al., 2010). The MEG signal is an average over a large population of neurons, and hence detection of population level high gamma responses requires precise (within a few ms) phase synchrony across these populations (Hämäläinen et al., 1993). However, the cortical sources which MEG would otherwise be sensitive to rarely phase synchronize across a large population at these high gamma ranges, leading to poor neural SNR for these high gamma ranges (reflected in our results by the small correlation values between actual responses and model predictions). The implications of this for our study are twofold. Firstly, conclusions regarding the intrinsic properties of high gamma responses to speech are limited by these methodological constraints on the MEG signal. Our results only show that there are significant cortical responses at ∼80 Hz, but do not rule out higher frequency cortical responses, or subcortical responses, that may be buried in poor SNR. Conversely, however, it is somewhat surprising that using such a simple experimental paradigm, with short duration continuous natural speech, it is possible to reliably detect such MEG responses using a TRF model.

### 4.2 MEG sensitivity to deep sources

Gradiometer-based MEG is physically constrained to be less sensitive to deep structures, typically resulting in such subcortical MEG responses being up to 100 times weaker than cortical responses at equivalent current strengths (Attal et al., 2007; Hillebrand and Barnes, 2002). Several source localization techniques have been proposed to correct for this inherent bias towards cortical sources (Dale et al., 2000; Krishnaswamy et al., 2017; Pascual-Marqui, 2002). Some studies were able to resolve MEG responses to the hippocampus (Cornwell et al., 2012), amygdala (Balderston et al., 2014; Cornwell et al., 2008; Dumas et al., 2013) and thalamus (Roux et al., 2013). Prior work has also been done using MEG for measuring brainstem responses (Coffey et al., 2016; Parkkonen et al., 2009). These studies show that MEG can be used to localize sources in subcortical areas given a large number of repetitions or specialized experimental paradigms. However, some of these studies used magnetometer-based MEG, which is more sensitive to deep sources than gradiometer based MEG (Lopes da Silva and van Rotterdam, 2005). In addition, resolving several sources with MEG is more complicated than localizing an isolated source due to the non-unique nature of distributed inverse solutions (Lütkenhöner, 2003). In our study, we used such a distributed source localization and a short experimental paradigm (without many stimulus repetitions) and found responses dominated by cortical sources. Simulation results suggested that the small amount of activation associated with the brainstem is more easily explained as an artifact of source localization leakage from cortical sources. On the other hand, although these results do not identify responses from subcortical regions, this does not imply at all that such responses are absent in the auditory system. Brainstem responses to speech have been detected using EEG (Maddox and Lee, 2018), and it is entirely possible that high gamma subcortical responses to speech may also be detected in MEG by other experiments with higher SNR, different analysis methods or MEG systems that are more sensitive to deep sources.

### 4.3 Cortical FFRs and high gamma TRFs

The high gamma TRF is not directly analogous to the FFR because, among other reasons, it is not an average over several repetitions of simple stimuli, whereas the TRF is a weighted average over longer time. However, the TRFs measured here indeed show a measure of high gamma time-locking that can be compared to the FFR. Cortical FFRs to repeated single speech syllables have been measured in MEG (Coffey et al., 2016) and EEG (Bidelman, 2018; Coffey et al., 2017b). Our work shows that cortical TRFs contain significant responses up to 62 ms, comparable to the long-lasting explanatory power of the auditory cortex ROI in Coffey et al., 2016. These TRFs are also predominantly from auditory cortex, centered around Heschl’s gyrus, and right lateralized similar to the MEG FFR (Coffey et al., 2016). However, some studies have demonstrated that the contribution of cortical sources to the FFR as measured with EEG is weaker than when measured with MEG (Ross et al., 2020), and rapidly decreases for harmonics above 100 Hz (Bidelman, 2018). In fact, while subcortical FFR is measurable with EEG for harmonics up to 1000 Hz, there were no cortical contributions to the FFR above 150 Hz (Bidelman, 2018). Unsurprisingly our results confirm that the cortical sources dominate the MEG response at frequencies near 100 Hz.

### 4.4 Comparison of responses to the envelope modulation vs. the carrier

The subcortical FFR is typically analyzed by averaging across stimulus presentations of opposite polarity, which results in responses driven mainly by the stimulus envelope and other even-order nonlinearities (Lerud et al., 2014). However, studies have also analyzed the FFR by subtracting the responses to stimulus presentations of opposite polarity (Aiken and Picton, 2008), which is driven mainly by the carrier and odd-order nonlinearities. Hence both the envelope and the carrier modulate responses across the auditory pathway. Unlike FFR analysis, the TRF analysis used in this study is well suited to disentangle the contributions of different features of the stimulus to the neural response, since it allows each stimulus representation to directly compete to explain the response variance. Our results found significant time-locked high gamma cortical responses to continuous speech for both envelope modulation and carrier, but these responses were predominantly driven by the envelope modulation over the carrier. This could be related to the perceptual phenomenon that modulation of the speech spectrum above 300 Hz is more behaviorally relevant for speech understanding, and more resistant to background noise, than the carrier below 200 Hz (Assmann and Summerfield, 2004). Slow evoked responses in auditory cortex are also sensitive to fine-structure acoustic features such as pitch and timbre (Roberts et al., 2000), and the auditory cortical response to the slowly varying envelope of speech is likewise modulated by the spectrotemporal fine structure of the stimulus (Ding et al., 2014).

### 4.5 High gamma TRF is driven by low pitch segments of the speech

The TRF response oscillates with a peak frequency of approximately 80 Hz, and is well time-locked to the segments of speech where the pitch is below 100 Hz. Cortical auditory phase locked responses to simple sounds have been measured using MEG (Coffey et al., 2016; Hertrich et al., 2004; Schoonhoven et al., 2003) at frequencies of up to 111 Hz. For continuous speech stimuli, such phase locked responses could reflect a cortical mechanism that represents complex speech features such as modulations in vowel formants, using fluctuations in the fundamental frequency domain of natural speech. Our pitch analysis showed that the response strongly locks to pitch frequencies below 100 Hz, but not above 100 Hz. This agrees with other studies that show a bias in cortical phase-locking towards lower frequencies (Bidelman, 2018; Ross et al., 2000; Schoonhoven et al., 2003).

### 4.6 Right lateralization of responses

The TRF model prediction accuracy was significantly right lateralized in younger subjects. The lack of significant right lateralization among older subjects may not indicate an age-related lateralization difference, but rather a lack of statistical power, since the lateralization was not significantly different across age groups. However, similar lateralization differences across age groups have been found for 80 Hz ASSR (Goossens et al., 2016). Stronger responses in the right auditory cortex have been observed for ASSR using EEG (Ross et al., 2005) and MEG (Hertrich et al., 2004) as well as in cortical FFRs using MEG (Coffey et al., 2016). This agrees with prior studies showing that right auditory cortex is specialized for early tonal processing and pitch resolution (Cha et al., 2016; Hyde et al., 2008; Zatorre, 1988). Both this right hemispheric bias, and the relatively short peak latency of 40 ms of our TRFs suggest that these cortical high gamma responses are due to early auditory processing of acoustic periodicity. However, some studies have also suggested that increased cortical folding in left auditory cortex could lead to a cancellation of MEG signals in the left hemisphere, which could lead to a similar right-ward bias in the absence of functional lateralization (Shaw et al., 2013).

### 4.7 Absence of age-related differences

Temporal precision and synchronized activity decreases in the auditory system with age and is characterized by age related differences in both subcortical and cortical responses. Older adults have subcortical FFR responses with smaller amplitudes, longer latencies and reduced phase coherence, which could be due to an excitation-inhibition imbalance or a lack of neural synchrony (Hornickel et al., 2012). In a surprising reversal, MEG and EEG studies have revealed that older adults have larger slow (< ∼10 Hz) cortical responses than younger adults (Alain et al., 2014; Bidelman et al., 2014; Herrmann et al., 2016), that result in better prediction accuracy for reverse correlation methods (Decruy et al., 2019; Presacco et al., 2016a, 2016b). Animal studies suggest that this opposite effect could be due to cortical compensatory central mechanisms (Chambers et al., 2016; Salvi et al., 2017) or lack of inhibition (Caspary et al., 2008; Villers-Sidani et al., 2010). Another possibility is the recruitment of additional neural areas for redundant processing (Brodbeck et al., 2018b; Peelle et al., 2010). Contrary to both these cases, we found no significant age-related differences in high gamma cortical responses, although this might be due to a lack of statistical power (see Ross et al., 2020). An ANOVA with factors TRF frequency and age suggested that the difference in low frequency responses among older and younger adults is not preserved for high gamma responses (see Appendix). These results suggest that high gamma cortical responses do not show a clear difference with age. The high gamma MEG TRF reflects fine-grained time-locked neural activity, like a subcortical FFR, but arising from cortical areas. It is possible that older adults’ exaggerated responses in cortical areas and the lack of neural synchrony at high frequencies (in subcortical FFRs) affect their high gamma MEG responses in opposite directions and obscure what would otherwise be detectable age-related differences.

### 4.8 Neural mechanisms for the MEG high gamma response

Given that MEG records the aggregate response over a large population of neurons, the specific origins of high gamma time-locked responses are not readily apparent. It is possible that the high gamma TRF reflects the effects of several processing stages along the auditory pathway, similar to the FFR. Electrocorticography (ECoG) studies have seen cortical phase-locked activity at these high rates (Nourski et al., 2014; Steinschneider et al., 2013). However, cortical phase-locking at the individual neuron level drastically reduces with increasing frequency (Lu et al., 2001), and hence cortical neurons may not be the sole contributor to these high gamma responses.

Such phase locked auditory activity is compatible with the spiking output of the Medial Geniculate Body (MGB) (Miller et al., 2002), which provides input to early auditory cortical areas. The MEG signal is dominantly driven by dendritic currents (that give rise to the Local Field Potential) (Hämäläinen et al., 1993), and hence these high gamma responses may be due to the inputs from the MGB into auditory cortex. Prior work has shown that auditory cortex is able to transiently time-lock to continuous acoustic features with surprisingly high temporal precision of the order of milliseconds (Elhilali et al., 2004). Time-locked inputs from MGB may provide a neural substrate for such precise transient temporal locking to stimulus features. Direct correspondences with age-related changes in thalamus from animal work are limited (Caspary and Llano, 2019), and hence it is unclear if time-locked high gamma spiking activity in MGB animal models would be similar across age. However, invasive neural recordings could help to disentangle the opposite effects of aging in the brainstem and the cortex seen with MEG and EEG, leading to a better understanding of time-locked responses in the aging auditory pathway.

## 5. Conclusion

In this study, we found high gamma time-locked responses to continuous speech, using MEG, that localized to auditory cortex, occurred with a peak latency of approximately 40 ms, and were stronger in the right hemisphere. We showed that TRF analysis could be used to reliably separate the contributions of several stimulus features to this response. The response function showed oscillations at approximately 80 Hz, predominantly driven by the envelope modulations during the segments of the speech where the pitch is below 100 Hz. Such high gamma time-locked responses may originate from the thalamic inputs to cortical neurons. These responses can be reliably detected in MEG using natural speech stimuli even of short duration, allowing TRF analysis to be employed to investigate auditory processing of speech from acoustics to semantics under several stimulus conditions in the same experiment. Furthermore, there were no significant age-related differences in these high gamma responses, unlike in both the low frequency cortical TRFs or the subcortical FFRs. Hence both the neural origin and the frequency domain must be considered when investigating age-related changes in the auditory system.

## Acknowledgements

Funding: This work was supported by the National Institute on Deafness and Other Communication Disorders (R01 DC-014085); the National Institute of Aging (P01 AG-055365); and the National Science Foundation (SMA1734892).

The identification of specific products or scientific instrumentation is considered an integral part of the scientific endeavor and does not constitute endorsement or implied endorsement on the part of the authors, DoD, or any component agency. The views expressed in this article are those of the authors and do not reflect the official policy of the Department of Army/Navy/Air Force, Department of Defense, or U.S. Government.

## Appendix

### Simulation of spatial spread of distributed source localization

Distributed neural source localization methods for MEG, such as MNE, result in a substantial amount of spatial spread. In order to characterize this spread, a dipole was simulated on Heschl’s gyrus perpendicular to the pial surface of the ‘fsaverage’ brain using the ‘ico-4’ surface source space. The dipole was then projected to sensor space, and MNE source localization with dSPM was performed to project it back onto the volume source space (see Fig A1). The peak of the activity shows a broad spread around Heschl’s gyrus but also some small activity in other parts of temporal lobe and even in the brainstem. This supports the claim that high gamma responses seen at the brainstem in our study are attributable to leakage from cortical areas.

**Fig. A1.**
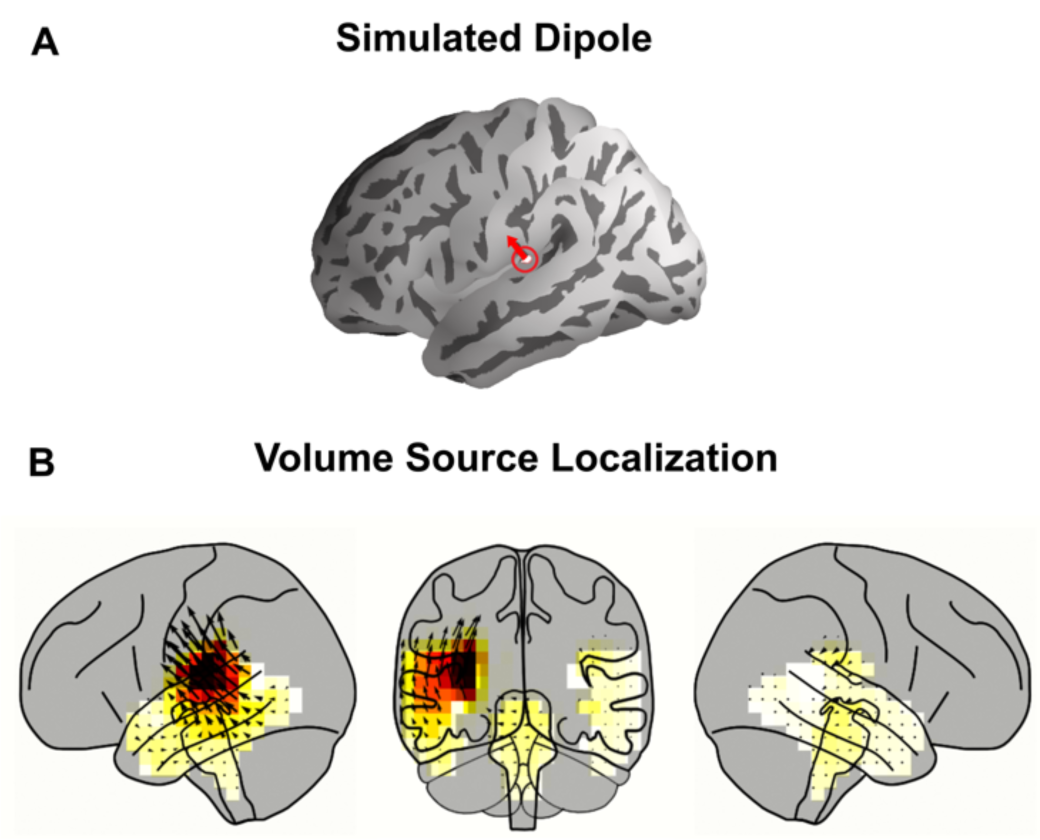
Simulation of Spatial Spread of Volume Source Localization. **A**. One dipole in Heschl’s gyrus was simulated. **B**. Volume Source Localization of the dipole after it was projected to sensor space. The cortical and subcortical ROIs are shown and artifactual leakage is seen in the brainstem voxels.

### Surface source space TRF methods and results

Cortical surface source space estimation was performed using the ‘ico-4’ source space, which consists of a fourfold icosahedral subdivision of the white matter surface of cortex with dipoles oriented normal to the surface. The ‘aparc’ parcellation was used to select dipoles in the temporal lobe for further analysis. In this surface source space analysis, current dipoles have a fixed orientation normal to the surface, and hence the TRF consists only of signed scalar amplitude variations with time. The Pearson correlation between the actual and predicted neural response was used as a measure of prediction accuracy for each neural source. For statistical tests, the TRFs and the correlation values were first rectified and then spatially smoothed using a Gaussian window with a standard deviation of 5 mm. The rectified, smoothed TRF of the true model was compared to the average of that of the three noise models using the same one tailed test with paired sample *t*-values and the TFCE procedure outlined in Methods.

Lateralization tests were performed to check for hemispheric asymmetry. The correlation values at each neural source in both left and right hemisphere were morphed onto the right hemisphere of the ‘fsaverage_sym’ brain as described in Brodbeck et al. (2018a). This brain model is symmetric in left and right hemispheres, allowing for comparisons between corresponding neural sources in both hemispheres. As before, these correlation coefficients were spatially smoothed using the same Gaussian window. After morphing, the correlation values of the average noise model were subtracted from that of the true model and a two-tailed test with paired sample *t*-values and TFCE was used to assess for significant differences in each of the corresponding left and right current dipoles.

TRFs were estimated using the cortical surface source space for neural sources in the temporal lobe, using both the envelope modulation and the carrier predictors in a competing model. Both predictors were time-shifted to generate noise models. All surface space results were similar to volume source space results. The prediction accuracies and TRFs are shown in Fig. A2, Fig. A3. The prediction accuracies were right lateralized but only in younger subjects (*t*_*max*_ = 4.6, *p* = 0.008). The TRFs showed a significant response in the range of 19–67 ms for the envelope modulation and 23–57 ms for the carrier. The envelope modulation TRF was stronger than the carrier TRF using the same tests as in the volume source space (younger *t*_*max*_ = 5.27, *p* < 0.001; older *t*_*max*_ = 3.46, *p* = 0.03). There were no age-related differences in surface source space analyses (prediction accuracy *t*_*max*_ = 2.41, *t*_*min*_ = –2.99, *p* > 0.29; maximum amplitude of envelope modulation TRF *t*_*max*_ = 2.40, *t*_*min*_ = –2.47, *p* > 0.68; maximum amplitude of carrier TRF *t*_*max*_ = 1.79, *t*_*min*_ = –3.07, *p* > 0.54).

In addition, low frequency TRFs were also estimated to compare age-related differences in both frequency domains. The stimulus representation for this model was the Hilbert envelope of the speech waveform filtered at 1–10 Hz with a logarithmic nonlinearity applied. The MEG data was also filtered at 1–10 Hz and TRFs were estimated using the surface source space. The resulting TRFs were as expected from prior work (Brodbeck et al., 2018b), with older subjects showing significantly higher reconstruction accuracies (*t*_*max*_ = 0.93, *t*_*min*_ = –3.45, *p* = 0.022). The increase in model prediction accuracies above the noise, for the high frequency TRF and the low frequency TRF were averaged across neural sources per subject, and a TRF frequency by age ANOVA was performed. Results indicated a significant interaction of TRF frequency x age (*F*_*1,38*_ = 6.46, *p* = 0.015) and significant main effects of TRF frequency (*F*_*1,38*_ = 216.58, *p* < 0.001) and age (*F*_*1,38*_ = 4.83, *p* = 0.034). This suggests that age-related changes are not consistent across low and high frequency responses, in further agreement with all the above results.

**Fig. A2.**
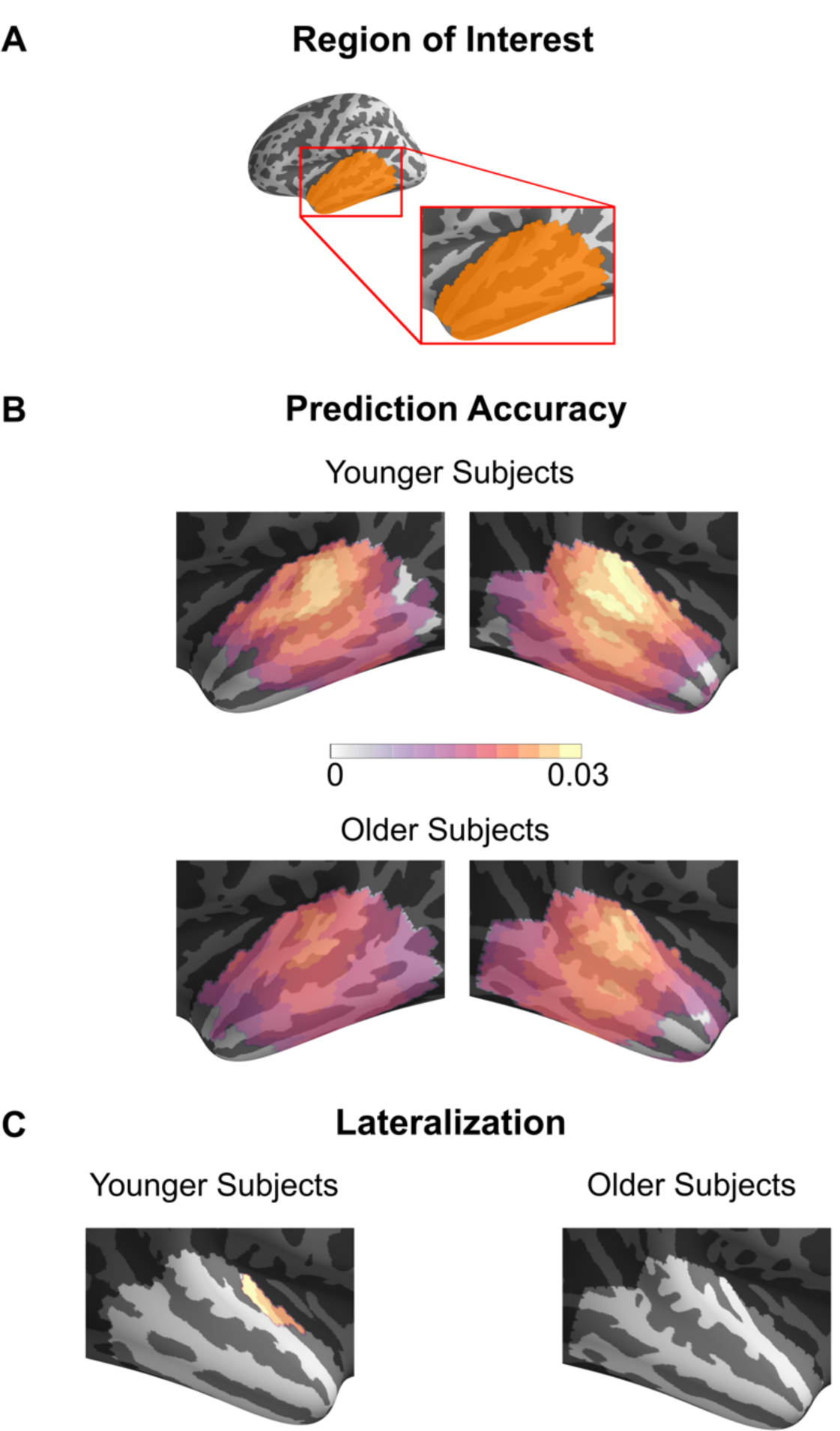
Prediction Accuracy of Surface Source Space TRFs. Pearson correlation coefficients between the actual and predicted response using the TRF model for each source in the surface source space ROI averaged across subjects are shown on an inflated brain. Only the voxels showing a significant increase in prediction accuracy over the noise model are plotted. Although most neural sources are significantly predictive, the prediction accuracy is larger in areas near core auditory cortex. A region in auditory cortex is significantly more predictive in the right hemisphere than the left, but only in younger subjects.

**Fig. A3.**
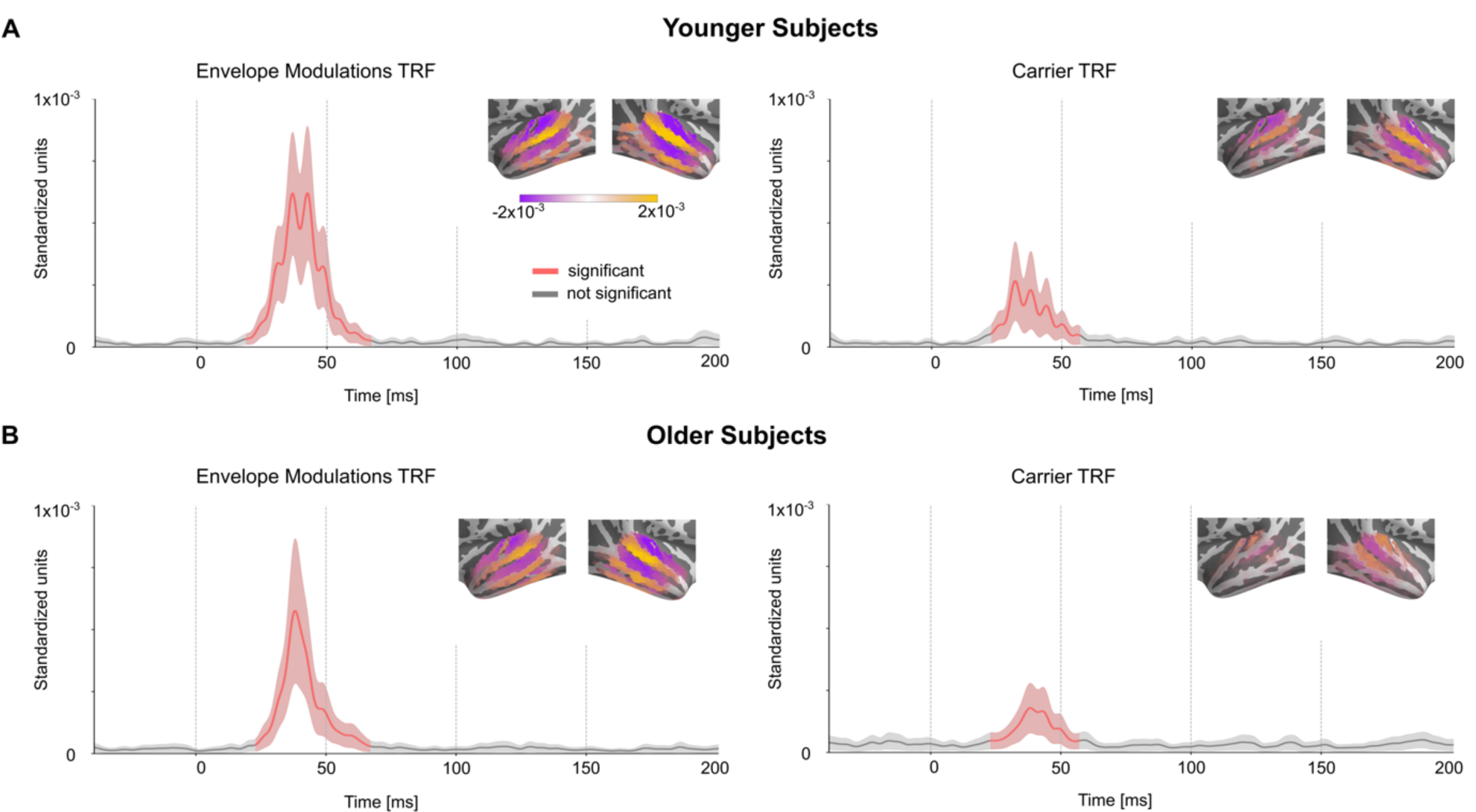
Surface Source Space TRFs. The amplitude of the TRFs for the competing model for both predictors averaged across neural sources and masked by significance against the noise model. Mean ± (standard error) across subjects is shown. The distribution of current dipoles in the temporal lobe ROI at the peak of the response is shown as an inset. Unlike the volume source space, the surface source space comprises of current dipoles with fixed orientation normal to the cortical surface. The signed magnitudes of these fixed direction dipoles are plotted on the surface, allowing for positive (orange) and negative (purple) values for outward and inward directions.

## References

Aiken, S.J., Picton, T.W., 2008. Envelope and spectral frequency-following responses to vowel sounds. Hear. Res. 245, 35–47. https://doi.org/10.1016/j.heares.2008.08.004

Alain, C., Roye, A., Salloum, C., 2014. Effects of age-related hearing loss and background noise on neuromagnetic activity from auditory cortex. Front. Syst. Neurosci. 8. https://doi.org/10.3389/fnsys.2014.00008

Anderson, S., Parbery-Clark, A., White-Schwoch, T., Kraus, N., 2012. Aging Affects Neural Precision of Speech Encoding. J. Neurosci. 32, 14156–14164. https://doi.org/10.1523/JNEUROSCI.2176-12.2012

Assmann, P., Summerfield, Q., 2004. The Perception of Speech Under Adverse Conditions, in: Greenberg, S., Ainsworth, W.A., Popper, A.N., Fay, R.R. (Eds.), Speech Processing in the Auditory System, Springer Handbook of Auditory Research. Springer, New York, NY, pp. 231–308. https://doi.org/10.1007/0-387-21575-1_5

Attal, Y., Bhattacharjee, M., Yelnik, J., Cottereau, B., Lefevre, J., Okada, Y., Bardinet, E., Chupin, M., Baillet, S., 2007. Modeling and Detecting Deep Brain Activity with MEG EEG, in: 2007 29th Annual International Conference of the IEEE Engineering in Medicine and Biology Society. Presented at the 2007 29th Annual International Conference of the IEEE Engineering in Medicine and Biology Society, pp. 4937–4940. https://doi.org/10.1109/IEMBS.2007.4353448

Baillet, S., 2017. Magnetoencephalography for brain electrophysiology and imaging. Nat. Neurosci. 20, 327–339. https://doi.org/10.1038/nn.4504

Balderston, N.L., Schultz, D.H., Baillet, S., Helmstetter, F.J., 2014. Rapid Amygdala Responses during Trace Fear Conditioning without Awareness. PLOS ONE 9, e96803. https://doi.org/10.1371/journal.pone.0096803

Basu, M., Krishnan, A., Weber-Fox, C., 2010. Brainstem correlates of temporal auditory processing in children with specific language impairment: Brainstem correlates of temporal processing. Dev. Sci. 13, 77–91. https://doi.org/10.1111/j.1467-7687.2009.00849.x

Bidelman, G.M., 2018. Subcortical sources dominate the neuroelectric auditory frequency-following response to speech. NeuroImage 175, 56–69. https://doi.org/10.1016/j.neuroimage.2018.03.060

Bidelman, G.M., 2015. Multichannel recordings of the human brainstem frequency-following response: Scalp topography, source generators, and distinctions from the transient ABR. Hear. Res. 323, 68–80. https://doi.org/10.1016/j.heares.2015.01.011

Bidelman, G.M., Villafuerte, J.W., Moreno, S., Alain, C., 2014. Age-related changes in the subcortical–cortical encoding and categorical perception of speech. Neurobiol. Aging 35, 2526–2540. https://doi.org/10.1016/j.neurobiolaging.2014.05.006

Boersma, P., 1993. Accurate short-term analysis of the fundamental frequency and the harmonics-to-noise ratio of a sampled sound. Proceedings of the Institute of Phonetic Sciences 97–110.

Boersma, P., Weenick, D., 2018. Praat: doing phonetics by computer [Computer program]. Version 6.0.43 [WWW Document]. URL http://www.praat.org (accessed 9.8.18).

Brodbeck, C., Brooks, T.L., Das, P., Reddigari, S., 2019. christianbrodbeck/Eelbrain: 0.30. Zenodo. https://doi.org/10.5281/zenodo.2653785

Brodbeck, C., Hong, L.E., Simon, J.Z., 2018a. Rapid Transformation from Auditory to Linguistic Representations of Continuous Speech. Curr. Biol. 28, 3976-3983.e5. https://doi.org/10.1016/j.cub.2018.10.042

Brodbeck, C., Presacco, A., Anderson, S., Simon, J.Z., 2018b. Over-Representation of Speech in Older Adults Originates from Early Response in Higher Order Auditory Cortex. Acta Acust. United Acust. 104, 774–777. https://doi.org/10.3813/AAA.919221

Brodbeck, C., Presacco, A., Simon, J.Z., 2018c. Neural source dynamics of brain responses to continuous stimuli: Speech processing from acoustics to comprehension. NeuroImage 172, 162–174. https://doi.org/10.1016/j.neuroimage.2018.01.042

Caspary, D.M., Ling, L., Turner, J.G., Hughes, L.F., 2008. Inhibitory neurotransmission, plasticity and aging in the mammalian central auditory system. J. Exp. Biol. 211, 1781–1791. https://doi.org/10.1242/jeb.013581

Caspary, D.M., Llano, D.A., 2019. Aging Processes in the Subcortical Auditory System, in: Kandler, K. (Ed.), The Oxford Handbook of the Auditory Brainstem. Oxford University Press, pp. 638–680. https://doi.org/10.1093/oxfordhb/9780190849061.013.16

Cha, K., Zatorre, R.J., Schönwiesner, M., 2016. Frequency Selectivity of Voxel-by-Voxel Functional Connectivity in Human Auditory Cortex. Cereb. Cortex N. Y. NY 26, 211–224. https://doi.org/10.1093/cercor/bhu193

Chambers, A.R., Resnik, J., Yuan, Y., Whitton, J.P., Edge, A.S., Liberman, M.C., Polley, D.B., 2016. Central Gain Restores Auditory Processing following Near-Complete Cochlear Denervation. Neuron 89, 867–879. https://doi.org/10.1016/j.neuron.2015.12.041

Coffey, E.B.J., Chepesiuk, A.M.P., Herholz, S.C., Baillet, S., Zatorre, R.J., 2017a. Neural Correlates of Early Sound Encoding and their Relationship to Speech-in-Noise Perception. Front. Neurosci. 11, 479. https://doi.org/10.3389/fnins.2017.00479

Coffey, E.B.J., Herholz, S.C., Chepesiuk, A.M.P., Baillet, S., Zatorre, R.J., 2016. Cortical contributions to the auditory frequency-following response revealed by MEG. Nat. Commun. 7, 11070. https://doi.org/10.1038/ncomms11070

Coffey, E.B.J., Musacchia, G., Zatorre, R.J., 2017b. Cortical Correlates of the Auditory Frequency-Following and Onset Responses: EEG and fMRI Evidence. J. Neurosci. 37, 830–838. https://doi.org/10.1523/JNEUROSCI.1265-16.2016

Cornwell, B.R., Arkin, N., Overstreet, C., Carver, F.W., Grillon, C., 2012. Distinct contributions of human hippocampal theta to spatial cognition and anxiety. Hippocampus 22, 1848–1859. https://doi.org/10.1002/hipo.22019

Cornwell, B.R., Carver, F.W., Coppola, R., Johnson, L., Alvarez, R., Grillon, C., 2008. Evoked amygdala responses to negative faces revealed by adaptive MEG beamformers. Brain Res. 1244, 103–112. https://doi.org/10.1016/j.brainres.2008.09.068

Dale, A.M., Liu, A.K., Fischl, B.R., Buckner, R.L., Belliveau, J.W., Lewine, J.D., Halgren, E., 2000. Dynamic Statistical Parametric Mapping: Combining fMRI and MEG for High-Resolution Imaging of Cortical Activity. Neuron 26, 55–67. https://doi.org/10.1016/S0896-6273(00)81138-1

David, S.V., Mesgarani, N., Shamma, S.A., 2007. Estimating sparse spectro-temporal receptive fields with natural stimuli. Netw. Bristol Engl. 18, 191–212. https://doi.org/10.1080/09548980701609235

de Cheveigné, A., Simon, J.Z., 2008. Sensor noise suppression. J. Neurosci. Methods 168, 195–202. https://doi.org/10.1016/j.jneumeth.2007.09.012

de Cheveigné, A., Simon, J.Z., 2007. Denoising based on Time-Shift PCA. J. Neurosci. Methods 165, 297–305. https://doi.org/10.1016/j.jneumeth.2007.06.003

Decruy, L., Vanthornhout, J., Francart, T., 2019. Evidence for enhanced neural tracking of the speech envelope underlying age-related speech-in-noise difficulties. J. Neurophysiol. 122, 601–615. https://doi.org/10.1152/jn.00687.2018

Ding, N., Chatterjee, M., Simon, J.Z., 2014. Robust cortical entrainment to the speech envelope relies on the spectro-temporal fine structure. NeuroImage 88, 41–46. https://doi.org/10.1016/j.neuroimage.2013.10.054

Ding, N., Simon, J.Z., 2012. Emergence of neural encoding of auditory objects while listening to competing speakers. Proc. Natl. Acad. Sci. 109, 11854–11859. https://doi.org/10.1073/pnas.1205381109

Dumas, T., Dubal, S., Attal, Y., Chupin, M., Jouvent, R., Morel, S., George, N., 2013. MEG Evidence for Dynamic Amygdala Modulations by Gaze and Facial Emotions. PLoS ONE 8. https://doi.org/10.1371/journal.pone.0074145

Elhilali, M., Fritz, J.B., Klein, D.J., Simon, J.Z., Shamma, S.A., 2004. Dynamics of Precise Spike Timing in Primary Auditory Cortex. J. Neurosci. 24, 1159–1172. https://doi.org/10.1523/JNEUROSCI.3825-03.2004

Fischl, B., 2012. FreeSurfer. NeuroImage 62, 774–781. https://doi.org/10.1016/j.neuroimage.2012.01.021

Forte, A.E., Etard, O., Reichenbach, T., 2017. The human auditory brainstem response to running speech reveals a subcortical mechanism for selective attention. eLife 6, e27203. https://doi.org/10.7554/eLife.27203

Goossens, T., Vercammen, C., Wouters, J., Wieringen, A. van, 2016. Aging Affects Neural Synchronization to Speech-Related Acoustic Modulations. Front. Aging Neurosci. 8. https://doi.org/10.3389/fnagi.2016.00133

Gordon-Salant, S., Yeni-Komshian, G.H., Fitzgibbons, P.J., Barrett, J., 2006. Age-related differences in identification and discrimination of temporal cues in speech segments. J. Acoust. Soc. Am. 119, 2455–2466. https://doi.org/10.1121/1.2171527

Gramfort, A., 2013. MEG and EEG data analysis with MNE-Python. Front. Neurosci. 7. https://doi.org/10.3389/fnins.2013.00267

Gramfort, A., Luessi, M., Larson, E., Engemann, D.A., Strohmeier, D., Brodbeck, C., Parkkonen, L., Hämäläinen, M.S., 2014. MNE software for processing MEG and EEG data. NeuroImage 86, 446–460. https://doi.org/10.1016/j.neuroimage.2013.10.027

Hämäläinen, M., Hari, R., Ilmoniemi, R.J., Knuutila, J., Lounasmaa, O.V., 1993. Magnetoencephalography---theory, instrumentation, and applications to noninvasive studies of the working human brain. Rev. Mod. Phys. 65, 413–497. https://doi.org/10.1103/RevModPhys.65.413

Hansen, P.C., Kringelbach, M.L., Salmelin, R. (Eds.), 2010. MEG: an introduction to methods. Oxford University Press, New York.

Hartmann, T., Weisz, N., 2019. Auditory cortical generators of the Frequency Following Response are modulated by intermodal attention. NeuroImage 203, 116185. https://doi.org/10.1016/j.neuroimage.2019.116185

He, N., Mills, J.H., Ahlstrom, J.B., Dubno, J.R., 2008. Age-related differences in the temporal modulation transfer function with pure-tone carriers. J. Acoust. Soc. Am. 124, 3841–3849. https://doi.org/10.1121/1.2998779

Herrmann, B., Henry, M.J., Johnsrude, I.S., Obleser, J., 2016. Altered temporal dynamics of neural adaptation in the aging human auditory cortex. Neurobiol. Aging 45, 10–22. https://doi.org/10.1016/j.neurobiolaging.2016.05.006

Hertrich, I., Dietrich, S., Trouvain, J., Moos, A., Ackermann, H., 2012. Magnetic brain activity phase-locked to the envelope, the syllable onsets, and the fundamental frequency of a perceived speech signal. Psychophysiology 49, 322–334. https://doi.org/10.1111/j.1469-8986.2011.01314.x

Hertrich, I., Mathiak, K., Lutzenberger, W., Ackermann, H., 2004. Transient and phase-locked evoked magnetic fields in response to periodic acoustic signals. Neuroreport 15, 1687–1690. https://doi.org/10.1097/01.wnr.0000134930.04561.b2

Hillebrand, A., Barnes, G.R., 2002. A Quantitative Assessment of the Sensitivity of Whole-Head MEG to Activity in the Adult Human Cortex. NeuroImage 16, 638–650. https://doi.org/10.1006/nimg.2002.1102

Hopkins, K., Moore, B.C.J., 2011. The effects of age and cochlear hearing loss on temporal fine structure sensitivity, frequency selectivity, and speech reception in noise. J. Acoust. Soc. Am. 130, 334–349. https://doi.org/10.1121/1.3585848

Hornickel, J., Anderson, S., Skoe, E., Yi, H.-G., Kraus, N., 2012. Subcortical representation of speech fine structure relates to reading ability: NeuroReport 23, 6–9. https://doi.org/10.1097/WNR.0b013e32834d2ffd

Hyde, K.L., Peretz, I., Zatorre, R.J., 2008. Evidence for the role of the right auditory cortex in fine pitch resolution. Neuropsychologia 46, 632–639. https://doi.org/10.1016/j.neuropsychologia.2007.09.004

Kraus, N., Anderson, S., White-Schwoch, T., Fay, R.R., Popper, A.N., 2017. The Frequency-Following Response: A Window into Human Communication. Springer.

Krishnan, A., Xu, Y., Gandour, J.T., Cariani, P.A., 2004. Human frequency-following response: representation of pitch contours in Chinese tones. Hear. Res. 189, 1–12. https://doi.org/10.1016/S0378-5955(03)00402-7

Krishnaswamy, P., Obregon-Henao, G., Ahveninen, J., Khan, S., Babadi, B., Iglesias, J.E., Hämäläinen, M.S., Purdon, P.L., 2017. Sparsity enables estimation of both subcortical and cortical activity from MEG and EEG. Proc. Natl. Acad. Sci. U. S. A. 114, E10465–E10474. https://doi.org/10.1073/pnas.1705414114

Kulasingham, J., 2019a. High Frequency Cortical Processing of Continuous Speech in Younger and Older Listeners - Dataset. UMD DRUM. https://doi.org/10.13016/33pk-ltqh

Kulasingham, J., 2019b. High Frequency TRF: Code [WWW Document]. URL https://github.com/jpkulasingham/highfreqTRF (accessed 12.19.19).

Lalor, E.C., Power, A.J., Reilly, R.B., Foxe, J.J., 2009. Resolving Precise Temporal Processing Properties of the Auditory System Using Continuous Stimuli. J. Neurophysiol. 102, 349–359. https://doi.org/10.1152/jn.90896.2008

Lerud, K.D., Almonte, F.V., Kim, J.C., Large, E.W., 2014. Mode-locking neurodynamics predict human auditory brainstem responses to musical intervals. Hear. Res. 308, 41–49. https://doi.org/10.1016/j.heares.2013.09.010

Lopes da Silva, F.H., van Rotterdam, A., 2005. Biophysical aspects of EEG and Magnetoencephalographic generation, in: Electroencephalography, Basic Principles, Clinical Applications and Related Fields. Philadelphia: Lippincott Williams & Wilkins, pp. 1165 – 1198.

Lu, T., Liang, L., Wang, X., 2001. Temporal and rate representations of time-varying signals in the auditory cortex of awake primates. Nat. Neurosci. 4, 1131–1138. https://doi.org/10.1038/nn737

Lütkenhöner, B., 2003. Magnetoencephalography and its Achilles’ heel. J. Physiol.-Paris 97, 641–658. https://doi.org/10.1016/j.jphysparis.2004.01.020

Maddox, R.K., Lee, A.K.C., 2018. Auditory Brainstem Responses to Continuous Natural Speech in Human Listeners. eNeuro 5. https://doi.org/10.1523/ENEURO.0441-17.2018

Miller, L.M., Escabí, M.A., Read, H.L., Schreiner, C.E., 2002. Spectrotemporal Receptive Fields in the Lemniscal Auditory Thalamus and Cortex. J. Neurophysiol. 87, 516–527. https://doi.org/10.1152/jn.00395.2001

Muthukumaraswamy, S., 2013. High-frequency brain activity and muscle artifacts in MEG/EEG: A review and recommendations. Front. Hum. Neurosci. 7. https://doi.org/10.3389/fnhum.2013.00138

Nichols, T.E., Holmes, A.P., 2002. Nonparametric permutation tests for functional neuroimaging: A primer with examples. Hum. Brain Mapp. 15, 1–25. https://doi.org/10.1002/hbm.1058

Nourski, K.V., Steinschneider, M., McMurray, B., Kovach, C.K., Oya, H., Kawasaki, H., Howard, M.A., 2014. Functional organization of human auditory cortex: Investigation of response latencies through direct recordings. NeuroImage 101, 598–609. https://doi.org/10.1016/j.neuroimage.2014.07.004

Oldfield, R.C., 1971. The assessment and analysis of handedness: The Edinburgh inventory. Neuropsychologia 9, 97–113. https://doi.org/10.1016/0028-3932(71)90067-4

Parkkonen, L., Fujiki, N., Mäkelä, J.P., 2009. Sources of auditory brainstem responses revisited: Contribution by magnetoencephalography. Hum. Brain Mapp. 30, 1772–1782. https://doi.org/10.1002/hbm.20788

Pascual-Marqui, R.D., 2002. Standardized low-resolution brain electromagnetic tomography (sLORETA): technical details. Methods Find. Exp. Clin. Pharmacol. 24 Suppl D, 5–12.

Peelle, J.E., Gross, J., Davis, M.H., 2013. Phase-Locked Responses to Speech in Human Auditory Cortex are Enhanced During Comprehension. Cereb. Cortex 23, 1378–1387. https://doi.org/10.1093/cercor/bhs118

Peelle, J.E., Troiani, V., Wingfield, A., Grossman, M., 2010. Neural Processing during Older Adults’ Comprehension of Spoken Sentences: Age Differences in Resource Allocation and Connectivity. Cereb. Cortex N. Y. NY 20, 773–782. https://doi.org/10.1093/cercor/bhp142

Peelle, J.E., Wingfield, A., 2016. The Neural Consequences of Age-Related Hearing Loss. Trends Neurosci. 39, 486–497. https://doi.org/10.1016/j.tins.2016.05.001

Presacco, A., Jenkins, K., Lieberman, R., Anderson, S., 2015. Effects of Aging on the Encoding of Dynamic and Static Components of Speech. Ear Hear. 36, e352–e363. https://doi.org/10.1097/AUD.0000000000000193

Presacco, A., Simon, J.Z., Anderson, S., 2016a. Evidence of degraded representation of speech in noise, in the aging midbrain and cortex. J. Neurophysiol. 116, 2346–2355. https://doi.org/10.1152/jn.00372.2016

Presacco, A., Simon, J.Z., Anderson, S., 2016b. Effect of informational content of noise on speech representation in the aging midbrain and cortex. J. Neurophysiol. 116, 2356–2367. https://doi.org/10.1152/jn.00373.2016

Puschmann, S., Baillet, S., Zatorre, R.J., 2019. Musicians at the Cocktail Party: Neural Substrates of Musical Training During Selective Listening in Multispeaker Situations. Cereb. Cortex 29, 3253–3265. https://doi.org/10.1093/cercor/bhy193

Roberts, T.P.L., Ferrari, P., Stufflebeam, S.M., Poeppel, D., 2000. Latency of the Auditory Evoked Neuromagnetic Field Components: Stimulus Dependence and Insights Toward Perception. J. Clin. Neurophysiol. 17, 114–129.

Rosen, S., 1992. Temporal information in speech: acoustic, auditory and linguistic aspects. Philos. Trans. R. Soc. Lond. B. Biol. Sci. 336, 367–373. https://doi.org/10.1098/rstb.1992.0070

Ross, B., Borgmann, C., Draganova, R., Roberts, L.E., Pantev, C., 2000. A high-precision magnetoencephalographic study of human auditory steady-state responses to amplitude-modulated tones. J. Acoust. Soc. Am. 108, 679–691. https://doi.org/10.1121/1.429600

Ross, B., Herdman, A.T., Pantev, C., 2005. Right Hemispheric Laterality of Human 40 Hz Auditory Steady-state Responses. Cereb. Cortex 15, 2029–2039. https://doi.org/10.1093/cercor/bhi078

Ross, B., Tremblay, K.L., Alain, C., 2020. Simultaneous EEG and MEG recordings reveal vocal pitch elicited cortical gamma oscillations in young and older adults. NeuroImage 204, 116253. https://doi.org/10.1016/j.neuroimage.2019.116253

Roux, F., Wibral, M., Singer, W., Aru, J., Uhlhaas, P.J., 2013. The Phase of Thalamic Alpha Activity Modulates Cortical Gamma-Band Activity: Evidence from Resting-State MEG Recordings. J. Neurosci. 33, 17827–17835. https://doi.org/10.1523/JNEUROSCI.5778-12.2013

Salvi, R., Sun, W., Ding, D., Chen, G.-D., Lobarinas, E., Wang, J., Radziwon, K., Auerbach, B.D., 2017. Inner Hair Cell Loss Disrupts Hearing and Cochlear Function Leading to Sensory Deprivation and Enhanced Central Auditory Gain. Front. Neurosci. 10. https://doi.org/10.3389/fnins.2016.00621

Schoonhoven, R., Boden, C.J.R., Verbunt, J.P.A., de Munck, J.C., 2003. A whole head MEG study of the amplitude-modulation-following response: phase coherence, group delay and dipole source analysis. Clin. Neurophysiol. 114, 2096–2106. https://doi.org/10.1016/S1388-2457(03)00200-1

Shaw, M.E., Hämäläinen, M.S., Gutschalk, A., 2013. How anatomical asymmetry of human auditory cortex can lead to a rightward bias in auditory evoked fields. NeuroImage 74, 22–29. https://doi.org/10.1016/j.neuroimage.2013.02.002

Smith, J.C., Marsh, J.T., Brown, W.S., 1975. Far-field recorded frequency-following responses: Evidence for the locus of brainstem sources. Electroencephalogr. Clin. Neurophysiol. 39, 465–472. https://doi.org/10.1016/0013-4694(75)90047-4

Smith, J.C., Marsh, J.T., Greenberg, S., Brown, W.S., 1978. Human auditory frequency-following responses to a missing fundamental. Science 201, 639–641. https://doi.org/10.1126/science.675250

Smith, S.M., Nichols, T.E., 2009. Threshold-free cluster enhancement: Addressing problems of smoothing, threshold dependence and localisation in cluster inference. NeuroImage 44, 83–98. https://doi.org/10.1016/j.neuroimage.2008.03.061

Steinschneider, M., Nourski, K.V., Fishman, Y.I., 2013. Representation of speech in human auditory cortex: Is it special? Hear. Res. 305. https://doi.org/10.1016/j.heares.2013.05.013

Villers-Sidani, E. de, Alzghoul, L., Zhou, X., Simpson, K.L., Lin, R.C.S., Merzenich, M.M., 2010. Recovery of functional and structural age-related changes in the rat primary auditory cortex with operant training. Proc. Natl. Acad. Sci. 107, 13900–13905. https://doi.org/10.1073/pnas.1007885107

Yang, X., Wang, K., Shamma, S.A., 1992. Auditory representations of acoustic signals. IEEE Trans. Inf. Theory 38, 824–839. https://doi.org/10.1109/18.119739

Yellamsetty, A., Bidelman, G.M., 2019. Brainstem correlates of concurrent speech identification in adverse listening conditions. Brain Res. 1714, 182–192. https://doi.org/10.1016/j.brainres.2019.02.025

Zan, P., Presacco, A., Anderson, S., Simon, J.Z., 2019. Mutual information analysis of neural representations of speech in noise in the aging midbrain. J. Neurophysiol. 122, 2372–2387. https://doi.org/10.1152/jn.00270.2019

Zatorre, R.J., 1988. Pitch perception of complex tones and human temporal-lobe function. J. Acoust. Soc. Am. 84, 566–572. https://doi.org/10.1121/1.396834

